# Amyloid-beta precursor protein contributes to brain aging and learning decline

**DOI:** 10.1101/2024.10.11.617841

**Authors:** Dennis E.M. de Bakker, Mihaela Mihaljević, Kunal Gharat, Yasmin Richter, Sara Bagnoli, Frauke van Bebber, Lisa Adam, Farzana Shamim-Schulze, Oliver Ohlenschläger, Martin Bens, Emilio Cirri, Adam Antebi, Ivan Matić, Anja Schneider, Bettina Schmid, Alessandro Cellerino, Janine Kirstein, Dario Riccardo Valenzano

## Abstract

Brain aging is a key risk factor for many neurodegenerative diseases, yet its molecular and cellular mechanisms remain elusive. Amyloid-beta precursor protein (APP) is among the most studied proteins linked to brain pathology; however, its role in non-pathological brain aging remains poorly characterized. Here, we investigate the natural impact of APP on normal brain aging using the short-lived turquoise killifish (*Nothobranchius furzeri*), which exhibits rapid and spontaneous age-related decline. We found that a pyroglutamated APP derivative (APP_pE11_) accumulates intra-neuronally in an age-dependent manner, co-localizing with a marker of cell death. We found that intraneuronal APP_pE11_ is also present in brains from healthy elderly humans, suggesting deep evolutionary conservation. To determine APP’s role in spontaneous brain aging, we knock-out “amyloid precursor protein a” (*appa*) in killifish via CRISPR/Cas9. The lack of *appa* mitigated brain aging from a proteome-wide perspective, reduced age-related cell death and inflammation, and improved neuronal activity and learning capacity in aged individuals. Our findings show an ancestral and previously unrecognized role of amyloid-beta precursor protein in non-pathological brain aging, making it an ideal target for anti-aging interventions.

## Introduction

The aging brain undergoes several changes, including increased protein aggregation, inflammation, and neuronal degeneration, which is associated with a decline in cognitive capacity. These changes are often exacerbated during neurodegenerative diseases, which possibly represent alternative aging trajectories^1–3^. Amyloid-beta precursor protein (APP) and its derivatives, including amyloid beta (Aβ_1-42_) are highly studied peptides that aggregate in the brain of normal – i.e., non-pathological – aging individuals as well as Alzheimer’s disease (AD) patients^4,5^. Genetic and pharmacological induction of Aβ_1-42_ can drive neurodegenerative symptoms in worms^6,7^, flies^8,9^ and mice^10–12^, and monoclonal antibodies and vaccination strategies targeting Aβ_1-42_ are being developed to slow down cognitive decline in AD patients^13–18^. However, whether APP derivatives play a key role in normal aging and whether they can be targeted to reduce the degenerative brain phenotypes of the normal aging brain remains largely unexplored.

The investigation of the effect of spontaneous accumulation of APP and its derivatives during aging requires an animal model whose natural aging trajectory includes both APP(-derivative) accumulation, as well as neurodegenerative phenotypes. Therefore, we set out to investigate the APP dynamics in a naturally short-lived vertebrate, which displays robust brain aging phenotypes, the turquoise killifish (*Nothobranchius furzeri*)^3^. Turquoise killifish have a median lifespan ranging from three to eight months and rapidly develop a range of age-related phenotypes overlapping with human aging. These phenotypes include sarcopenia^19^, cancer susceptibility^20^, telomere attrition^21,22^, impaired regenerative capacity^23,24^, as well as a decrease in antibody diversity and increased immunosenescence^25–27^. Meanwhile, the killifish brain is subjected to age-related protein aggregation^28–33^, astrogliosis^34,35^, inflammation^24,28,35,36^, neuronal degeneration^29,37–39^ and reduced neurogenic potential^24,34,35^, which together might underlie a decreased learning capacity^39–41^. Since several brain aging phenotypes arise spontaneously, the turquoise killifish has been proposed as a promising model to study spontaneous age-dependent neurodegenerative processes ^3,42^.

Here, we investigate the role of amyloid-beta precursor protein (APP) in normal brain aging. Using transcriptomics, proteomics, and immunohistochemistry, we show that with age, the killifish brain becomes subjected to a robust inflammatory state accompanied by intracellular accumulation of a pyroglutamated APP-derivative, a post-translational modification associated with increased toxicity^43,44^. Finally, we show that knocking out the *appa* isoform of the amyloid-beta precursor genes mitigates age-related brain phenotypes.

## Results

### Amyloid-beta precursor protein is evolutionary conserved between humans and turquoise killifish and is expressed in the aged turquoise killifish brain

The turquoise killifish is a naturally short-lived species, reaching sexual maturity within 3 to 5 weeks post-hatching. In this study, we define killifish (strain ZMZ1001) as “young,” “aged,” and “old” based on their ages of 6 weeks, 6 months, and 9 months, respectively. These age categories correspond to survival rates of 100%, 89%, and 44% (**Figure 1a**). Old individuals display several visually apparent aging phenotypes, including spine curvature, craniofacial alterations, as well as melanocytotic tailfins in males (**Figure 1b**). The brain morphology of the turquoise killifish resembles that of a typical teleost (**Figure 1c**) and has been extensively described elsewhere^45^. The turquoise killifish genome contains two paralogs of the human amyloid precursor protein (APP), which encode the amyloid beta (Aβ) sequence, *appa* and *appb* (coding sequences provided in **Supplementary Table 1**). Both paralogs show a highly conserved Aβ_1-42_ sequence from human to killifish from the 11^th^ amino acid onwards (Aβ_11-42_), as well as conserved beta- and gamma-secretase cleavage sites (**Figure 1d**). Alphafold3 indicates similar 3D architecture for killifish (*appa* derived) and human (*APP* derived) Aβ_1-42_ and predicts that despite low conservation at the N’ side, killifish Aβ_1-42_ can effectively form multimers (**Extended Data Figure 1a-e**). To investigate whether APP derivatives are detectable in the aged turquoise killifish brain, we performed immunohistochemistry using two Aβ-targeting antibodies. Considering that the available antibodies are not specifically raised against the killifish Aβ peptide, together with the known risks of misinterpretation of non-specific signals in the aged brain^46,47^, we first addressed the binding specificity of the antibodies. To that end, we used the 4G8 antibody, targeting the Aβ_17-24_ epitope which is fully conserved between humans and killifish, and can bind full-length APP and various APP derivatives including Aβ. Additionally, we used the pE11 antibody, which targets N-terminally truncated and pyroglutamated Aβ, through recognition of the N-terminal pyroglutamate group and the first six amino acids after truncation (Aβ_pE11-16_) (**Figure 1d**). While 4G8 broadly marks full-length APP and its derivatives, pE11 specifically targets post-translationally modified Aβ (Aβ_pE11-40/42_) in human samples, enabling the tracking of cleaved and toxic Aβ rather than its precursors. We focused on the pE11 antibody over the pE3 antibody, as the pE11 target epitope is conserved between humans and killifish. Even though the 4G8 and pE11 antibodies bind different epitopes of the Aβ peptide, we found a strong co-localization between 4G8 and pE11 signal throughout the anterior rhombencephalon (AR) of aged (6-month-old) killifish (**Figure 1e**). To mitigate the possibility that both antibodies bind the same structure in an epitope-independent manner, we blocked the epitopes by incubating antibodies with synthetic Aβ_pE11-42_ peptide prior to the immunohistochemical staining. While incubation with albumin did not impact the staining pattern of pE11 or 4G8, the signal of both antibodies was almost completely abolished by the pre-incubation with Aβ_pE11-42_ peptide, indicating that both antibodies bind to their targets in the killifish brain through their epitope binding domains (**Extended Data Figure 2a,b**). Noteworthy, while pE11 antibody binding indicates the presence of an APP derivative which is cleaved and pyroglutamated at the 11^th^ amino acid of its Aβ sequence, it does not provide evidence for C’ cleavage of APP. Therefore, we hereafter refer to pE11-bound APP derivatives as APP_pE11_. Taken together, we conclude that the aged turquoise killifish brain contains APP derivatives, including APP_pE11_.

**Figure 1.**
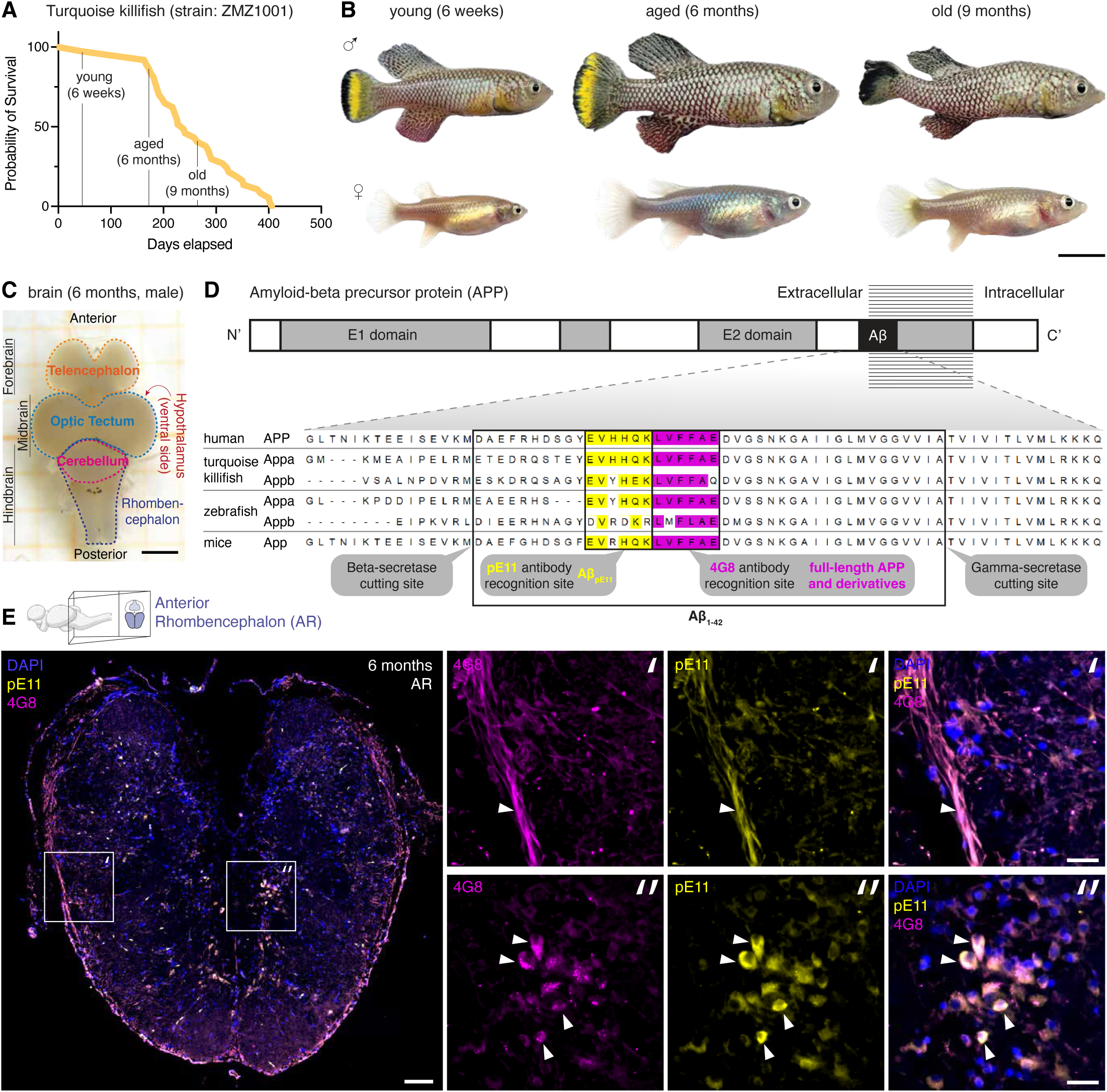
Amyloid-beta precursor protein is evolutionary conserved between turquoise killifish and humans and is expressed in the turquoise killifish brain. **(A)** Survival curve for the turquoise killifish strain ZMZ1001. **(B)** Representative pictures of male and female fish at different ages. Scalebar represents 1 cm. **(C)** Representative brain (6-month-old male) with annotated brain regions. Scalebar represents 1 mm. **(D)** Graphical representation of the amyloid beta precursor protein (APP) and amino acid sequences of APP precursors across four species. Amino acid residues are colored according to their physicochemical properties (ZAPPO): aliphatic/hydrophobic (residues ILVAM) - rose, aromatic (FWY) - orange, positive (KRH) - dark blue, negative (DE) - red, hydrophilic (STNQ) - light green, conformationally special (PG) - magenta, cysteine (C) - yellow. **(E)** Immunohistochemistry staining in the 6-month-old anterior rhombencephalon (AR) using the pE11 and 4G8 antibodies, targeting AβpE11_-42_ (yellow) and APP-derivatives (magenta), respectively. Arrowheads indicate pE11+/4G8+ signal. Scalebars represent 100 *µ*m in the overview and 25 *µ*m in the zoom-ins.

### Turquoise killifish develop age-related brain inflammation while expressing key genes involved in post-translational APP processing

To investigate the age-dependent mRNA expression levels of *appa* and *appb*, as well as other key genes involved in the biogenesis of APP derivatives, we performed whole brain RNA-sequencing on six young (6-week-old) and six aged (6-month-old) female turquoise killifish (**Supplementary Table 2**). Interestingly, genes downregulated with age indicate a strong decline in cellular proliferation (e.g. *mki67*), consistent with the reported loss of neurogenic potential^24,34,35^ (**Extended Data Figure S3a,b**). Genes upregulated with age (e.g., *stat1*, *irf9*, *rsad2*, *cmpk2*) suggest a strong age-related induction of inflammation (**Extended Data Figure 3c**), in agreement with previous findings^24,28,35,36^. To test a potential age-related inflammatory state in the turquoise killifish brain, we adopted markers of brain inflammation in the hypothalamus and optic tectum and tested them via immunohistochemistry by targeting Lymphocyte Cytosolic Protein 1 (LCP1), a pan-leukocyte marker employed to visualize microglia in the zebrafish and killifish brain^24,35,48^ (**Extended Data Figure 3d,e**). As LCP1 is an actin-binding protein, the staining pattern visualizes cytoskeletal architecture of brain leukocytes (**Extended Data Figure 3f**). We found a strong age-related increase in the surface area covered by LCP1, which validates the age-related inflammatory state suggested by the transcriptomics data (**Extended Data Figure 3g**). Next, to adress whether key genes involved in APP derivative biogenesis are actively transcribed in the killifish brain and would support a potential role for APP and it’s derivatives during killifish brain aging, we plotted their expression in young and aged brains (**Extended Data Figure 3h**). These genes include amyloid precursor proteins *appa*, *appb*, the APP family members *appl1*, *appl2*, the glutaminyl cyclase *qpctl* which pyroglutamates Aβ, the beta-secretase *bace1* and the family member *bace2* as well as the gamma-secretase subunits *psen1*, *psen2*, *ncstn* and *aph1b*. While all factors showed consistent expression in the turquoise killifish brain, no age-related changes were observed at the mRNA level. In summary, the aging killifish brain becomes subjected to a robust inflammatory state and expresses transcripts encoding key genes involved in the biogenesis of APP derivatives.

### pE11 signal accumulates across the rostro-caudal axis of the killifish brain

Next, we asked whether *appa* signal detected via 4G8 and pE11 (**Figure 1d,e**) shows any age-related changes. We found that the anterior rhombencephalon (AR) of young, 6-week-old fish, presents 4G8 signal covering a similar percentage of the tissue area as in the aged 6-month-old counterparts (**Figure 2a-c**), possibly reflecting the binding of 4G8 to full-length APP. In contrast, the young AR appears largely devoid of any pE11 signal, while pE11 signal was instead detected in the aged AR (**Figure 2d-f**). These findings suggest that while full-length APP and its derivatives might be present in the brain throughout killifish lifespan, APP_pE11_ accumulates only at a later age. To explore the full spatiotemporal extent of the accumulation of pyroglutamated Aβ, we characterized the pE11 signal in seven different brain regions along the rostro-caudal axis, at four different ages (**Figure 2g, Extended Data Figure 4-8**). While the pE11 signal remained nearly undetectable until three months of age, the pE11 covered surface area peaked at six months, which was consistent across all investigated brain regions. At nine months of age, the pE11 covered surface area was decreased compared to the six-month timepoint. To test for possible batch effect explaining signal changes between six and nine months, we repeated this experiment and confirmed the finding in an independent cohort (**Extended Data Figure 9**). The decline in pE11 covered surface area between 6- and 9-months of age potentially suggests an increase in clearance, or a reduced production of APP_pE11_. Alternatively, the reduction of pE11 between 6- and 9-months could be due to the death of pE11^+^ cells, and/or due to the selective disappearance of animals with high APP_pE11_ levels. Taken together, these results indicate that pyroglutamated APP derivatives accumulate spontaneously across the rostro-caudal axis between 3- and 6-months of age in the brain of the turquoise killifish.

**Figure 2.**
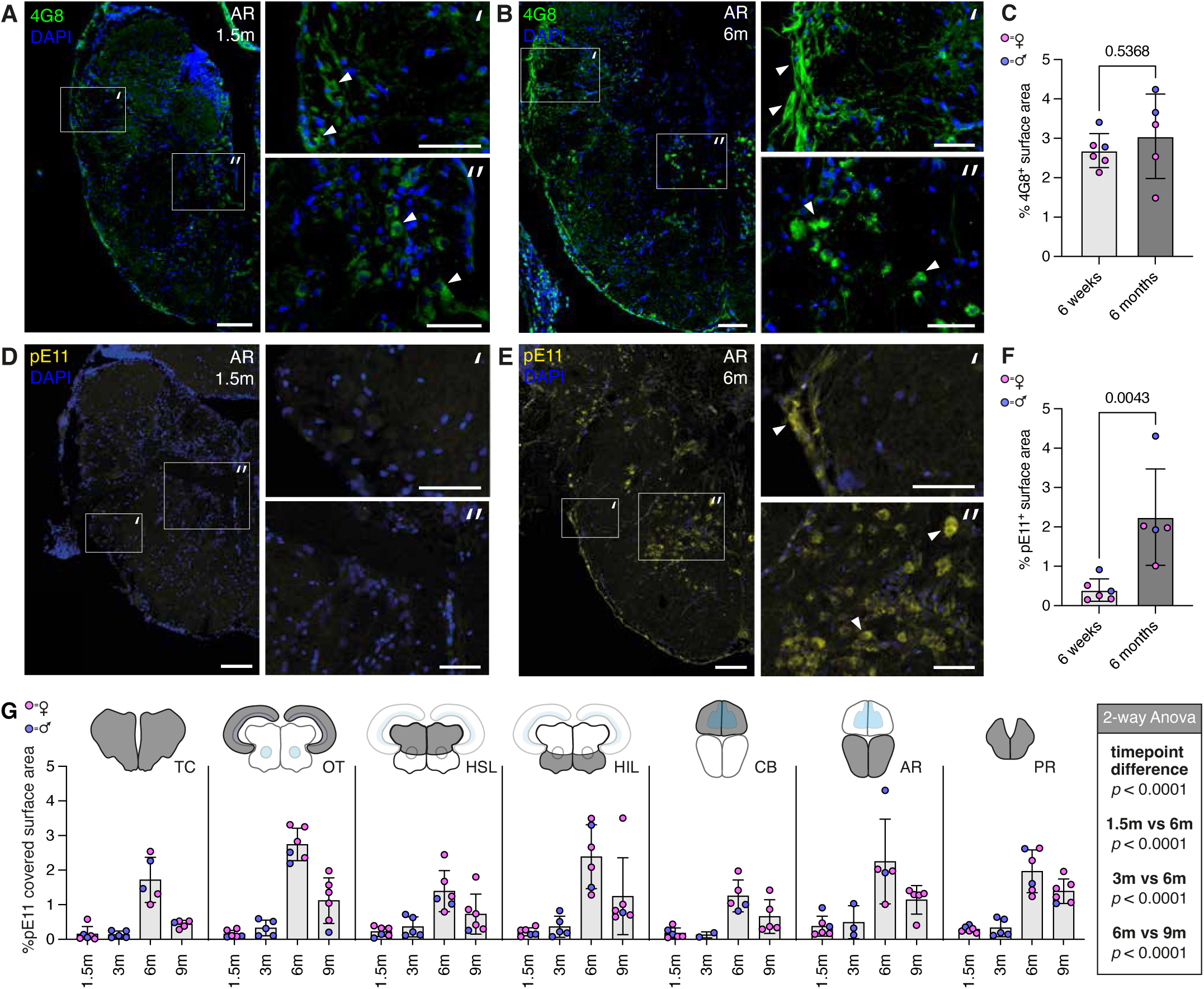
pE11 signal accumulates in an age-dependent manner across the rostro-caudal axis of the turquoise killifish brain. (**A, B**) Immunohistochemistry staining using the 4G8 antibody in the anterior rhombencephalon (AR) of young (6-week-old) and aged (6-month-old) killifish. (**C**) Quantification of the percentage of 4G8-covered surface area in 6-week-old and 6-month-old AR regions. (**D, E**) Immunohistochemistry staining using the pE11 antibody in the anterior rhomboencephalon (AR) of young (6-week-old) and aged (6-month-old) killifish. (**F**) Quantification of the percentage of pE11-covered surface area in 6-week-old and 6-month-old AR regions. (A, B, D, E) Scale bars represent 100 μm in the overview and 50 μm in the zoom-in. Arrowheads indicate positive signal. (C, F) Significance was calculated using a Mann-Whitney test. Error bars represent standard deviation. (**G**) Quantification of the percentage of pE11-covered surface area across seven different brain regions at four different time points, including the telencephalon (TC), optic tectum (OT), hypothalamus superior lobe (HSL), hypothalamus inferior lobe (HIL), cerebellum (CB), anterior rhombencephalon (AR) and posterior rhombencephalon (PR). Error bars represent standard deviation. Significances were calculated using a 2-way Anova test, including a multiple comparison test to investigate the difference between 1.5-months and 6-months, 3-months and 6-months as well as 6-months and 9-months.

### APP_pE11_ predominantly localizes intra-neuronally and colocalizes with TUNEL

In the human AD etiology, the APP derivative Aβ is typically known to localize in extracellular beta-sheet aggregates known as Aβ plaques. However, Aβ has also been described to localize intraneuronally^49–54^. We could not identify any large pE11^+^ or 4G8^+^ extracellular signals indicative of the formation of extracellular amyloid plaques in the aged killifish brain. Instead, we found that 4G8 and pE11 signal mainly localized around nuclei, suggesting a predominantly intracellular localization (**Figure 1e, Extended Data Figure 4-8**). To investigate the potential intraneuronal localization of pE11 in the aged killifish brain, we co-labeled pE11 together with an antibody targeting neuronal nuclei (NeuN).

Indeed, pE11 localized directly around NeuN^+^ nuclei (**Extended Data Figure 10a**), indicating that APP_pE11_ predominantly localizes intra-neuronally. To address whether the intra-neuronal localization of APP_pE11_ is unique to the killifish or whether its localization is evolutionary conserved and also present in aged human brains, we performed pE11 staining on the brains of normal non-pathological (>80 years old) and AD diagnosed individuals (**Extended Data Figure 10b, 11, 12**). Strikingly, we detected intraneuronal pE11 signal in both the cortex and the hippocampus of normal aged human brains, as well as in the brains of Alzheimer’s patients. In addition, we detected 4G8 and pE11 signal in extracellular plaques of the Alzheimer’s disease brains, which is in line with previous reports^55^. These findings indicate that the occurrence of pyroglutamated APP derivatives is not limited to a specific human pathology, but instead is broadly present the in aged vertebrate brain in an intraneuronal fashion. Pyroglutamated Aβ has been shown to be exceedingly neurotoxic^43^. Therefore, to investigate whether pE11^+^ cells show signs of DNA double stranded breaks typically indicative of programmed cell death, we adopted TUNEL staining. TUNEL^+^ cells were nearly undetectable in young killifish brains but were clearly detectable in aged brains (**Extended Data Figure 13, 14a**). Importantly, in aged brains TUNEL signal colocalized with pE11 signal (**Extended Data Figure 14b**), which might suggest that spontaneous accumulation of APP_pE11_ could negatively affect neuronal survival in aged killifish. Indeed, we could detect TUNEL^+^/NeuN^+^ neurons throughout the aging killifish rhombencephalon (**Extended Data Figure 15**), in agreement with previous findings indicating that the aging killifish brain is subjected to substantial neuronal degeneration^29,37–39^.

### APP derivatives aggregate in an age-related manner

The aging killifish brain is characterized by substantial protein aggregation^28–33^. To determine whether this includes Aβ-associated beta-sheet aggregation, we conducted a thioflavin-T assay. This assay detects changes in the fluorescent emission profile of thioflavin-T upon its incorporation into beta-sheet aggregates^56^. Aged brain lysates yielded higher thioflavin-T signal compared to their younger counterparts, indicating an age-dependent increase in beta-sheets protein aggregates (**Extended Data Figure 16**). To directly test whether APP derivatives show signs of age-related aggregation, we performed a filter retardation assay, where protein lysates derived from killifish brains of several ages are applied to a filter which retains protein aggregates of a specific size determined by the pore size of the filter ^57^. Next, we stained the filter-bound aggregated protein with either the 4G8 or the pE11 antibody. We found that while 4G8 signal was enriched in an age-dependent manner (**Extended Data Figure 17a,b**), this was not the case for the detected pE11 signal (**Extended Data Figure 17c,d**). Corroborating the potential age-dependent aggregation of 4G8^+^/pE11^-^ peptides, we detected an age-dependent increase in intracellular 4G8^+^/pE11^-^ puncta in the killifish brain (**Extended Data Figure 18**), which may reflect intracellular aggregation of APP derivatives. Indeed, 4G8^+^ puncta showed a partial overlap with the Aggresome dye, which was previously employed to visualize intracellular protein aggregation in the killifish brain^30^ (**Extended Data Figure 19**). The aged human brain display a similar pattern between 4G8 and pE11 signals, as the 4G8 signal showed a granulated pattern in contrast to the pE11 staining (**Extended Data Figure 20**). Taken together, our results indicate that while the aged killifish brain is largely devoid of extracellular plaques, APP derivatives (4G8^+^/pE11^-^) aggregate in an age-dependent and intracellular manner.

### CRISPR/Cas9 mediated knock-out of *appa* results in reduction of APP derivatives

Given that intraneuronal APP_pE11_ signal as well as 4G8^+^ puncta mark the aged killifish and human brain, we asked whether APP derivatives contribute to brain aging. To test this hypothesis, we generated an amyloid-beta precursor protein a (*appa*) mutant killifish line using CRISPR/Cas9. Using two guide RNAs targeting the second exon of the *appa* gene, we introduced a 16bp deletion leading to a frameshift (**Figure 3a**). The induced frameshift did not lead to a premature stop-codon but led to a scrambled amino acid coding sequence, which prevented the translation of *appa* derived APP peptide after the second exon, including the amyloid beta coding sequence. In addition to *appa*, we also targeted its homolog, *appb*, for knock-out. While successfully inserting mutations in the *appb* gene, founders could not be raised to adulthood and were therefore excluded from the analysis ^58^. The loss of *appb* mutant carriers might indicate a role for *appb* in killifish development, which is in line with the described role for *appb* during early embryonic development in the zebrafish^59^. To investigate whether *appb* compensates the lack of *appa*-derived APP, we performed a Western blot using the 22C11 antibody targeting the full-length APP. 22C11 should target both wild-type APPA and APPB, as the epitope targeted by 22C11 is highly conserved in both homologs (16/16 amino acids) and might also detect killifish APLP2 (12/16). APLP1 is less conserved (9/16) and will likely not be detected by the 22C11 antibody. We observed a significant reduction (>50%) of APP levels in the killifish *appa* mutant brain lysates, indicating that the still functional *appb* could not compensate in full for the loss of *appa* derived APP (**Figure 3b,c**). The lack of *appa* in the knock-out fish line did not affect survival rates, nor did the *appa* mutants show specific signs of discomfort (**Figure 3d**). At 6-months of age, the reduced levels of APP led to a reduction in APP derivatives, including APP_pE11_, indicated by the observed reduction in 4G8 and pE11 immunoreactivity in both the anterior rhombencephalon and hypothalamus superior lobe (**Extended Data Figure 21, Figure 3e-g**). This effect was statistically investigated using a Bayesian model, as it allowed for the bespoke integration of observed slide-to-slide variation in signal intensity. The reduced immunoreactivity of both antibodies in the *appa* mutants further supports their specific binding to *appa* derived peptides. Residual antibody immunoreactivity likely stemmed from *appb* derived peptides. Taken together, the *appa* knock-out killifish line showed reduced levels of APP and its derivatives, including APP_pE11_.

**Figure 3.**
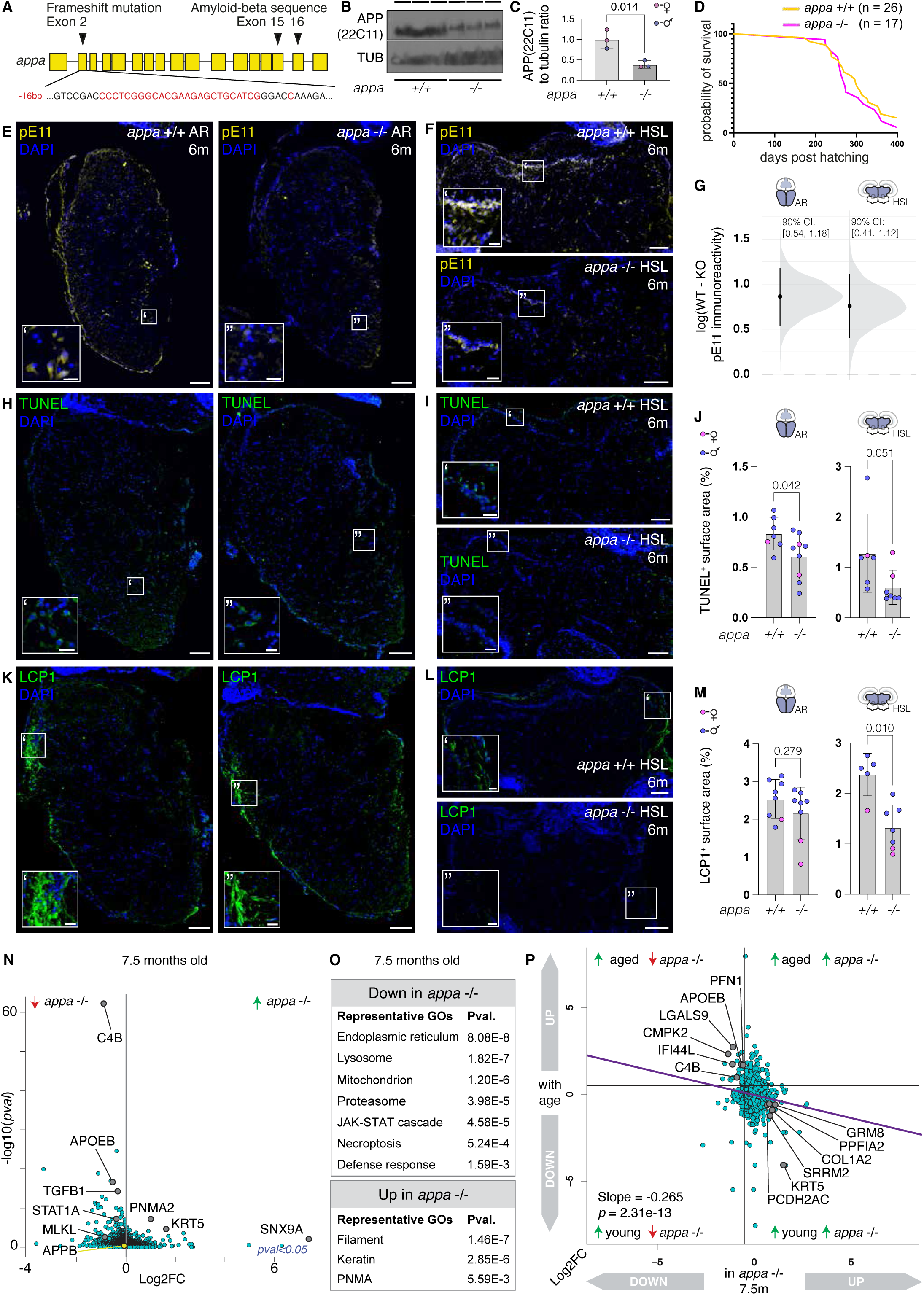
*appa* knock-out mitigates killifish brain aging. (**A**) Schematic representation of the killifish *appa* gene, indicating the mutation site on exon 2, as well as highlighting the location of the amyloid beta coding region in exons 15 and 16. (**B**) Western blot on turquoise killifish lysates from 6-month-old killifish using APP(22C11) and alpha-tubulin targeting antibodies. (**C**) Quantification of western blot results, depicting the APP(22C11) to tubulin ratio. Significance was calculated using a Mann-Whitney test. (**D**) Survival curve of a cohort of wild-type (yellow, n = 26) and *appa* -/- (magenta, n = 17) turquoise killifish of the ZMZ1001 strain. (**E, F**) Representative immunohistochemistry staining results using the pE11 antibody (yellow) in the (E) anterior rhombencephalon (AR) and (F) hypothalamus superior lobe (HSL) of wild-type and *appa* -/- brains. Scale bar represents 100 µm in the overviews and 20 µm in the magnified view. (**G**) Posterior estimates of the genotype effect (WT KO) on pE11 immunoreactivity were obtained using a Bayesian model. Black vertical bars represent the 90% confidence intervals (CIs).The results indicate higher pE11 immunoreactivity in *appa* wild-type (WT) brains compared to *appa* -/- (KO) brains in the anterior rhombencephalon (AR) and hypothalamus superior lobe (HSL) of 6-month-old killifish. (**H, I**) Representative TUNEL staining results (green) in the anterior rhomb-encephalon (AR) and hypothalamus superior lobe (HSL) of wild-type and *appa* -/- brains. (**J**) Quantification of the percentage of TUNEL-covered surface area in anterior rhomboencephalon (AR) and hypothalamus superior lobe (HSL) regions of 6-month-old individuals. Significance was c alculated using a Mann-Whitney test. Error bars represent standard deviation. (**K, L**) Representative immunohistochemistry staining results using the LCP1 antibody (green) in the anterior rhombencephalon (AR) and hypothalamus superior lobe (HSL) of wild-type and *appa* -/- brains. Scale bar represents 100 µm in the overviews and 20 µm in the magnified view. (**M**) Quantification of the percentage of LCP1-cov-ered surface area in anterior rhomboencephalon (AR) and hypothalamus superior lobe (HSL) regions of 6-month-old individuals. Significance was calculated using a Mann-Whitney test. Error bars represent standard deviation. (**N**) Volcano plot displaying differences in brain protein abundance between *appa* -/- mutants and their wild-type siblings (7.5-month-old, 3 males and 3 females per genotype). Differentially abundant proteins are highlighted, representing relevant biological processes. The vertical line indicates log2FC of 0. (**O**) Representative gene ontolo-gies produced from the differentially abundant proteins between 7.5-month-old *appa* -/- mutants and their wild-type siblings (pval < 0.05). (**P**) Proteome-wide comparison between the log2FC of proteins up or downregulated in 6-months-old brains compared to 7-weeks-old brains (y-axis), plotted against the log2FC describing the up and downregulation of proteins in the *appa* mutant compared to the same aged wild-type siblings at 7.5 months.

### Knock-out of *appa* mitigates brain aging

To investigate the effects of *appa* knock-out on specific aging phenotypes, we took a histological approach and assessed levels of cell death and inflammation. To visualize cell death in the *appa* mutants, we performed TUNEL staining in tissue sections of the anterior rhombencephalon and hypothalamus superior lobe. While TUNEL^+^ cells were consistently present in brains of wild-type sibling, *appa* mutant brains contained a lower TUNEL^+^ covered surface area in both investigated regions at as early as 6-months of age (**Figure 3h-j**). To further validate the potential mitigation of age-related inflammation, we performed an immunohistochemical staining using an antibody targeting Lymphocyte Cytosolic Protein 1 (LCP1) as previously described (**Extended Data Figure 3d-f**). The 6-months-old *appa* mutant brain displayed a significant reduction of LCP1 covered surface area in the hypothalamus superior lobe, but not in the anterior rhombencephalon (**Figure 3k-m**).

To investigate the effects of the knock-out of *appa* in an unbiased manner, we performed liquid chromatography mass spectrometry (LC-MS) on whole brain lysates (**Supplementary Table 3**). We used three males and three females from both *appa* mutants, as well their wild-type siblings, for each of the two age groups (6- and 7.5-month-old). We chose the 7.5 months time-point as it corresponds to the median lifespan of fish from the ZMZ1001 strain (**Figure 1a**), when behavioral changes become apparent and therefore corresponds to a critical timepoint to assess brain aging phenotypes related to cognition^39,41^. At both timepoints, *appa* derived peptides were undetectable in the *appa* mutants, while *appb* derived peptides (APPB) showed no significant changes (**Figure 3n, Extended Data Figure 22a**). At 6-months of age, the *appa* mutants displayed an upregulation of proteins involved in mitochondria (e.g., DHRS12, SLC25A3, NDUFV1), translation (e.g., RPS7, RPL27, RPS13), and DNAJ-chaperones (e.g., DNAJC5AA, DNAJA1L, DNAJB3, DNAJA3A), as well as a downregulation of amyloidosis associated factors Apolipoprotein A1 (APOA1) and beta-synuclein (SNCB) (**Figure 22a,b**). Compared to same-aged wild-type siblings, 7.5-month-old *appa* mutants showed an upregulation of keratins (e.g., KRT4, KRT5, KRT18) and a downregulation of proteins involved in the lysosome and proteasome, as well as proteins involved in inflammation related ontologies, such as the defense response (e.g., C4B, TGFB1) and the JAK-STAT cascade (e.g., STAT1A). In addition, we observed downregulation of proteins involved in necroptosis (e.g., MLKL, XIAP) which is suggestive of a reduction in programmed cell death (**Figure 3n,o**)^59^. Taken together, the LC-MS and histological findings indicate that the knock-out of *appa* mitigates spontaneous cell death and inflammation in the aging turquoise killifish brain.

To determine whether the changes in protein abundances upon *appa* knock-out represent a broader mitigation of brain aging, we investigated whether we could detect a proteome-wide shift towards a younger state in the *appa* mutants compared to their wild-type siblings. To this end, protein group abundances were compared between young (6-week-old) and aged (6-month-old) brains, as well as between *appa* mutant and wild-type sibling brains at two different ages (6 and 7.5 months). Subsequently, the log2 fold-change (log2FC) values from both the young-versus-old and *appa* mutant-versus-wild-type comparisons were plotted against one another. In 6-month-old *appa* mutants, the differentially abundant proteins did not correlate with age-related changes in protein level (**Figure 22c**). However, in 7.5-month-old *appa* mutants, we observed a highly significant negative correlation (slope = −0.265, *p* = 2.31e-13) indicating that proteins upregulated with age were downregulated in the mutants, and *vice versa* (**Figure 3P**). Proteins upregulated with age but downregulated in the *appa* mutants included proteins implicated in neurodegenerative diseases (e.g., APOEB, PFN1, LGALS9) and inflammation (e.g., IFI44L, C4B, CMPK2), while proteins downregulated with age and upregulated in the *appa* mutants included extracellular matrix components (e.g., COL1A2, KRT5) as well as proteins involved in neuronal function (e.g., GRM8, PCDH2AC, PPFIA2). These findings indicate that *appa* knock-out mitigated brain aging in a proteome-wide manner between 6- and 7.5 months of age, extending beyond the reduction of inflammation and cell death.

### Knock-out of *appa* mitigates age-related learning decline

Turquoise killifish display a spontaneously occurring decline in learning capacity with age, a *bona fide* measure of age-related cognitive decline^39,41^. To investigate whether *appa* contributes to non-pathological age-dependent learning deficiency, we developed an active avoidance test based on previous literature^39,60,61^. Each fish (28 *appa* mutants and 30 wild-type siblings) was individually tested in an arena consisting of two circular compartments, connected through a narrow opening (**Figure 4a**). After a 15-minute acclimatization period, fish were subjected to 40 consecutive cycles, where a conditionate stimulus (light), was coupled with a non-conditioned stimulus (air bubbles). Each one-minute cycle corresponded to 30 seconds of light exposure, with the latter 15 seconds combined with exposure to air bubbles, followed by a rest phase of 30 seconds. Untrained fish showed a preference to remain in the light, with 84% of the fish staying within the light during the first cycle of the experiment (**Figure 4b**). Due to the pairing of the light with an adverse non-conditioned stimulus (air bubbles) across the 40 cycles, the fish learn to avoid air bubbles once cued with light. Indeed, even 7.5-month-old wild-type fish (n = 30) become increasingly proficient at actively avoiding the light with 16% (first cycle) to 46% (last cycle) of the fish escaping the bubbles (**Figure 4b, Extended Data Figure 23, Supplementary Table 4**). However, the 7.5-month-old *appa* mutants (n = 28) outperform their age-matched wild-type siblings and show an increased rate of successful escapes after the 6^th^ cycle onwards, while reaching an average escape rate of 55% during the last cycle (**Figure 4b, Extended Data Figure 23**). Comparing the total amount of successes, as well as their top scores, defined as the maximum number of successes in a 10-cycle window, show higher (yet non statistically significant) learning performance in the *appa* -/- mutants (*p* = 0.073 and *p* = 0.265, respectively) (**Figure 4c,d**). Significant improvements in the *appa* -/- mutants were observed when comparing the speed at which their top scores were reached (**Figure 4e**) and when considering both top scores and the speed (i.e. the number of the learning cycle) at which the top score was reached simultaneously (**Figure 4f**). To gain insight into the biological significance of the learning improvements observed in the *appa* -/- mutants at 7.5-months of age, we performed the active avoidance test on an independent cohort of young killifish (6-week-old, n = 16) (**Extended Data Figure 24**). While young fish show a higher escape rate in the first 5 cycles of the experiment compared to all other non-young counterparts, the 7.5-month-old mutants performed not only better than age-matched controls, but similar to young fish for the remaining cycles. These results suggest that knock-out of *appa* can almost completely rescue the age-related learning deficit observed in turquoise killifish.

**Figure 4.**
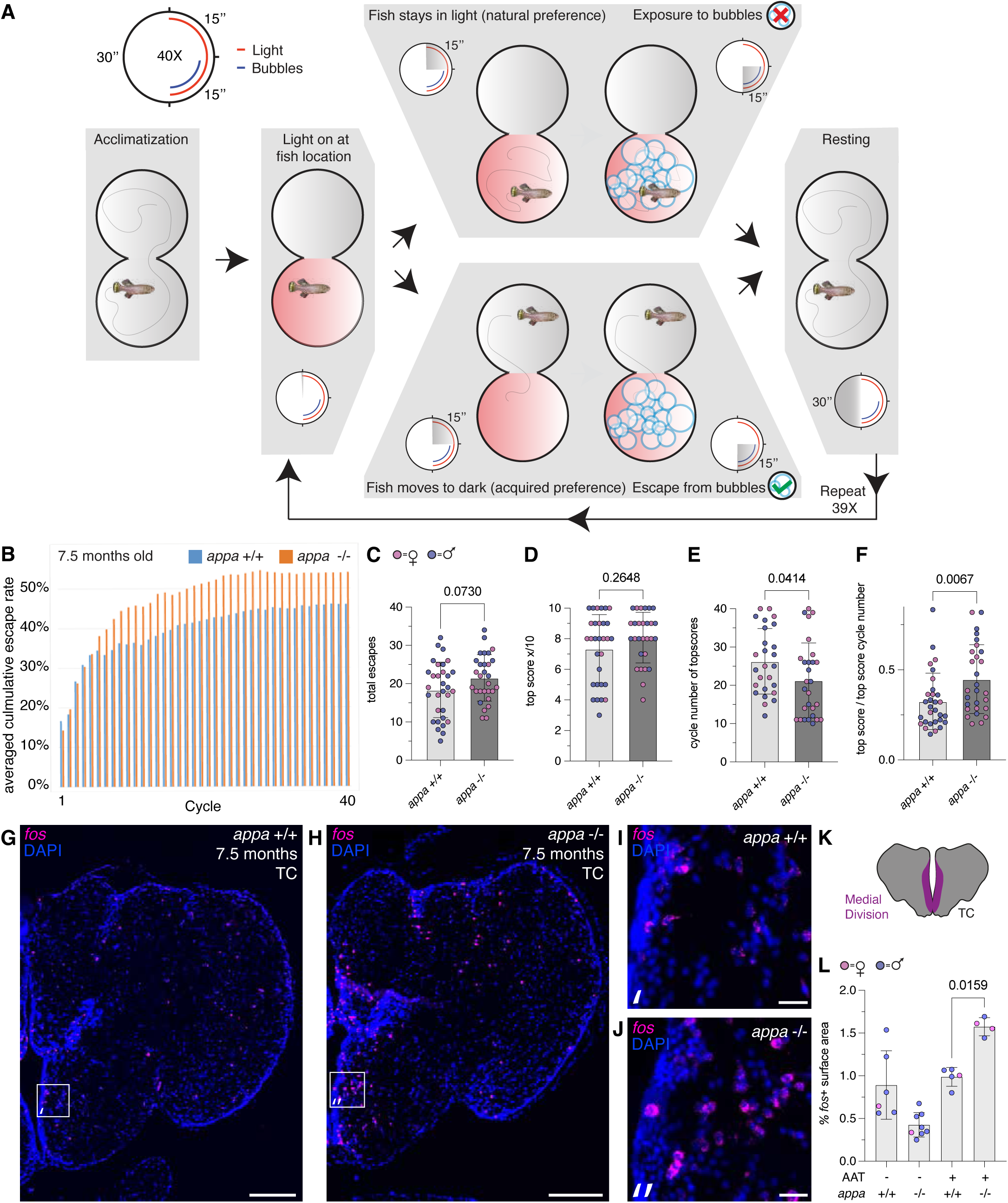
*appa* knock-out mitigates age-related learning deficit in turquoise killifish. **(A)** Experimental set-up of the performed active avoidance test (AAT). (**B**) Plot describing the averaged cumulative escape rate of 7.5-month-old *appa* -/- mutants (orange, n = 28 fish) and their wild-type siblings (blue, n = 30 fish) across 40 cycles. (**C-F**) Quantification of four behavioral traits, compared between 7.5-month-old *appa* -/- mutants and their wild-type siblings. These traits include (**C**) the total escape rate (n/40 cycles), (**D**) the top score of the fish within a window of ten consecutive cycles representing the extent of learning, (**E**) the cycle number at which top scores were reached (excluding non-learners (n=4) defined by total escapes < 10) representing the speed of learning, and (**F**) a combined value in which both the extent and speed of learning were considered (top scores / cycle number at which top scores were reached). Significance values were calculated through Mann-Whitney testing. (**G-J**) Hybridizing Chain Reaction (HCR) staining against *fos* transcripts (magenta), marking neurons involved in active learning within the telencephalon (TC) of *appa* mutants (**H,J**) and their wild-type siblings (**G,I**). Scale bars represent 200 µm in the overview and 20 µm in the zoom-ins. (**K**) Schematic representation of the medial division (magenta) within the telencephalon (TC). (**L**) Quantification of the percentage of *fos*^+^ surface area in the medial division of the telencephalon of 6-month-old *appa* -/- mutant fish and their wild-type siblings that have not been subjected to an active avoidance test (AAT -), as well as as 7.5-month-old *appa* -/- mutant fish and their wild-type siblings which were subjected to an active avoidance test (AAT +). Significance value was calculated through a Mann-Whitney test.

Newly learned behaviors (e.g. the negative association with red light) are encoded within a neuronal network which, upon activation, expresses the immediate response factor c-Fos^62,63^. To address whether the active avoidance test we employed led to the activation of such network, and whether the *appa* mutants show increased network activity compared to their wild-type siblings, we used Hybridization Chain Reaction (HCR) *in situ* mRNA detection. We used HCR to visualize *fos* transcripts in brains isolated immediately after the 40th cycle of the active avoidance task (**Figure 4g-j**). We specifically quantified the percentage surface area covered by *fos* transcripts in the medial zone of the telencephalon (**Figure 4k**), considering that this region has been previous implicated in learning performance of other fish species^64^. We found that the *appa* mutants that underwent the active avoidance test showed higher *fos* levels in the medial zone when compared to untested fish, as well as their wild-type siblings that did undergo the active avoidance test (**Figure 4l**). From these results, we conclude that the knock-out of *appa* rescued an age-induced learning deficit, which is associated with enhanced neuronal activity in the telencephalon.

## Discussion

In this study, we asked whether the detrimental effect of APP derivatives on the brain is limited to neurodegenerative diseases, or whether APP derivatives also impact normal, non-pathological brain aging. Although APP derivatives (e.g. Aβ_1-42_) has been shown to accumulate during normal aging^65^, convincing experimental proof of its causal effect remained absent. An important clue suggesting a potential causal effect of APP derivatives in the normal aging brain was provided by a previous study describing the cognitive fitness of aged human carriers of the Islandic APP variant (A673T). The A673T APP mutation leads to a reduction in β-Secretase 1 (BACE1)-mediated Aβ production^66^, which not only correlated with a reduced risk to develop Alzheimer’s disease (AD), but also with increased cognitive fitness in normal aging individuals^67^. In addition, a negative correlation was found between amyloid burden and cognitive performance in non-pathological centenarians^68^. Here, we provide the first experimental evidence showing that lowering APP levels in spontaneously aging vertebrates can reduce brain aging phenotypes and mitigate age-related cognitive decline.

To investigate the effect of APP during normal brain aging, we have employed the turquoise killifish. These annual teleosts evolved a compressed lifespan due to the genome-wide accumulation of risk alleles, largely driven by past population bottlenecks^69^. Therefore, the aging phenotypes observed in the turquoise killifish brain are most likely a consequence of their exceptionally high mutation load and have a polygenic basis. Due to their short lifespan, turquoise killifish represent an ideal model for functional studies investigating vertebrate aging. In this study, we recapitulate previously reported brain aging phenotypes of the turquoise killifish, including the aggregation of protein, inflammation, and cellular degeneration. Furthermore, we describe an additional brain aging phenotype, the intraneuronal accumulation of APP_pE11_. Notably, we found that intraneuronal APP_pE11_ not only characterizes the aging brain of the killifish but is also present in aged, non-diseased human brains. Intraneuronal APP_pE11_ had previously been associated with Alzheimer’s disease^70^ but, to the best of our knowledge, remained unexplored in the context of normal brain aging. While Aβ_pE3_ is expressed and studied in AD mouse models (e.g. 5xFAD)^71^, no animal data were available to date for the endogenous expression of APP_pE11_. Several mechanisms could explain intraneuronal APP_pE11_ accumulation during aging, including an age-dependent increase in APP endocytosis^72^ further processed in the *trans*-Golgi network^73,74,75^, or a decline in APP_pE11_ clearance efficiency.

To test the impact of APP during normal aging, we knocked out one the amyloid-beta precursor protein coding genes, *appa*. Both the histological and LC-MS results indicated that *appa* knock-out mitigated cellular death and inflammation in aged killifish brains. Furthermore, proteome-wide analysis indicated that the beneficial effects of *appa* knock-out are not restricted to inflammation and cell death but encompass a much broader, proteome-wide mitigation of brain aging. The beneficial effects of *appa* knock-out on brain aging in the turquoise killifish is in stark contrast with the negative effects observed in APP knock-out mice, which are viable but have altered neurogenesis and smaller brains^78^. In addition, aged APP knock-out mice display increased reactive gliosis in the neocortex and hippocampus and show reduced locomotor activity^79^, as well as reduced memory retention^80^. It has been suggested that both excessive amounts of Aβ, as well as complete depletion of Aβ, has negative effects on brain functionality^81^. As teleosts, turquoise killifish have been subjected to a whole genome duplication^82^ and hence, contain two APP paralogues. Therefore, while knockout of APP in mice resulted in a complete depletion of Aβ, knock-out of *appa* in killifish left the fish with *appb*, which might have yielded a concentration of Aβ high enough to allow typical brain development and function, but low enough to evade Aβ-mediated exacerbation of brain aging phenotypes.

While decades of research have focused on elucidating the effects of APP and its derivatives during Alzheimer’s disease, the effects of APP during normal aging remained largely unexplored. Here, we show that APP exacerbates brain aging in a short-lived teleost. These findings indicate that the detrimental effects of APP and its derivatives are not limited to a specific human pathology, but could instead broadly impact vertebrate brain aging. APP-targeting strategies, originally intended to mitigate neurodegenerative diseases, may therefore have broader potential as an anti-aging approach to preserve cognitive health throughout life.

## Methods

### Fish husbandry and maintenance

The turquoise killifish (ZMZ1001 strain) were housed in individual 2.8L tanks equipped with baffles, fry mesh and lids glass capillaries (Aquaneering, USA) connected to a system with circulating water. Fish received 12 h of light and 12 h of darkness per day. Water temperature was set to 28°C. Fish were fed with brine shrimp nauplii (*Artemia salina*) and blood worm larvae (*Chironomus spp.*) twice a day during the week and once a day during the weekend. Adult fish (6 weeks post hatching) were used for breeding purposes. One male and up to three females were housed in 9.5L breeding tanks equipped with baffles, fry mesh and lids glass capillaries (Aquaneering, USA). Fertilised eggs were regularly collected using sand boxes and then stored in a petridish with coconut fiber at 24°C. Three weeks prior to hatching, embryos were placed at 28°C. Furthermore, the husbandry and maintenance of the killifish used in this study were performed as part of the standard operating procedure the Leibniz institute on Aging, which was described in detail in a previous publication^83^. The experiments were carried out under permission of the following animal experimental licenses: FLI-21-016 (young behavioral tests), FLI-23-029 (appa KO versus wild-type behavioral tests), Par.11 J-003798 (Maintenance and Housing), FLI-22-203 (Genotyping), O-DV-23-27 (Organ harvest license).

### Protein alignment and Alphafold protein structure predictions

To determine how similar protein sequences of APPA and APPB from turquoise killifish are to protein sequences from humans (*H. sapiens*), mice (*M. musculus*) and zebrafish (*Danio rerio*), full-length peptide sequences were aligned using UniPro UGENE (RRID:SCR_005579). Multiple protein sequences were aligned with the MUSCLE default mode. The sequences of all protein variants from online database Ensembl (https://www.ensembl.org/index.html). Structure predictions of Aβ_1-42_ were performed using the AlphaFold3 web server with default options^84^. For graphical representations the program ChimeraX^85^ was employed.

### Brain processing for cryo-sectioning

Killifish were euthanized using rapid-chilling method. The whole head was first fixed in 4% paraformaldehyde overnight at room temperature. The next day brain was isolated, washed in 4% sucrose and incubated in 30% sucrose overnight at 4°C. On the last day, brain was transferred to an appropriate Tissue-Tek cryomold, immersed in the Tissue-Tek O.C.T. Compound and transferred to the dry ice to quickly freeze the tissue. Brains were stored at −80°C until the sectioning step. The tissue blocks were sectioned to 10 μm thin sections using cryostat machine, while following standard protocol for cryo-sectioning.

### Immunohistochemistry, imaging and quantification

Thawed slides were first washed in PEMTx solution (80 mM 98.5+% PIPES (1,4-Piperazin-bis-(ethansulfonacid), 5 mM EGTA, 1 mM MgCl_2_, 0.2% Triton X-100 in milliQ water, pH 7,5) for 10 min, in NH_4_Cl-PEMTx (50 mM of NH_4_Cl in PEMTx) for another 10 min, and then two more times in PEMTx. The slides were transferred and incubated in Sodium Citrate Solution (10 mM 99% Trisodium Citrate Dihydrate in RO H_2_O, pH 6) for 15 min at 85°C in the waterbath. Once cooled down to the room temperature, slides were washed two times for 10 min in PEMTx. The tissue was incubated in the blocking solution (10% Fetal Bovine Serum, 1% DMSO in PEMTx) for 1 hour at room temperature in a humidified chamber. Slides were later incubated with primary antibodies diluted in a blocking buffer at 4°C overnight in a humidified chamber. The following day, the slides were washed in PEMTx solution five times for 20 min (on a shaker). The slides were incubated in the secondary antibody (1:500) and DAPI (1:1000) diluted in a blocking buffer for 1 h at room temperature in a humidified chamber and in the dark. The primary antibody Rabbit Anti-Abeta-pE11 was combined with the secondary antibody Alexa Fluor® 647 Donkey anti-rabbit IgG (min. x-reactivity) antibody (BioLegend, USA, Cat# 406414, RRID: AB_2563202). The Rabbit Anti-GFAP antibody was combined with the secondary antibody Alexa Fluor 555 (Thermo Fisher Scientific, USA, Cat# A-21428, RRID: AB_2535849). The slides were washed in PEMTx solution five times for 10 min, then two times for 7 min in PEM solution (80 mM 98.5+% PIPES (1,4-Piperazin-bis-(ethansulfonacid), 5 mM EGTA, 1 mM MgCl_2_ in miliQ H_2_O) on the shaker and in the dark. Slides were mounted in the ProLong Gold antifade reagent and stored at 4°C in the dark. The slides were imaged using Zeiss Axio Scan.Z1 microscope with a Plan-Apochromat 20x/0.8 M27 objective, Axiocam 506m imaging device and 1x Camera Adapter. Imaging results were processed using ZEN 3.5 (Blue Edition) ZEN Digital Imaging for Light Microscopy (RRID:SCR_013672) to select images and Fiji (ImageJ2, RRID:SCR_002285) and Imaris (V10.2, 2024 June 26) to quantify positive signal. Positive signal was defined as the percentage of the surface area within a given brain region that was covered by staining. For each brain region and individual, three sections were analyzed, and mean values were calculated for presentation in the figures.

### Signal intensity measurements and Bayesian statistical analysis

Brains from *appa* wild-type and mutant fish were isolated and embedded in pairs. Sectioning at 10 µm thickness generated slides containing both wild-type and mutant tissue.

Immunohistochemistry was performed using the 4G8 and pE11 antibodies. Signal intensity was quantified as the mean pixel intensity across the tissue-covered area within defined brain regions (Figure 3g, Figure S21c). We quantified the signal intensity of 4G8 and pE11 in the *appa* mutants, instead of the signal covered surface area, as we observed that while the covered surface area remained relatively stable, the signal intensity appeared visibly reduced. For each brain region and individual, three stained sections were analyzed.

Imaging was carried out using a Zeiss Axio Scan.Z1 microscope equipped with a Plan-Apochromat 20×/0.8 M27 objective, an Axiocam 506m camera, and a 1× camera adapter. Image acquisition and signal quantification were performed with ZEN 3.12 software.

Because *appa* wild-type and mutant brains were embedded and sectioned together, we were able to observe slide-to-slide variability in average signal intensity, despite consistent staining and imaging protocols. To account for this, we applied a Bayesian model to estimate the genotype effect (wild-type – knockout) on immunoreactivity. This approach incorporated slide-to-slide variation as a covariate and yielded posterior distributions from which confidence intervals were derived.

### Immunohistochemistry on human brain samples

Human brain tissue samples of AD patients and aged-matched controls were obtained from the London Neurogenerative Diseases Brain Bank, King’s College London. For each individual, we used consecutive tissue sections of 10 um thickness in order to visualize the precise localization of the staining for 4G8 and pE11 in comparable areas.

Tissue sections deriving from 2 AD patients ( BBN-ID: BBN_9888, 80, M, and BBN_002.34142, 77, M) and 2 control individuals ( BBN-ID: BBN002.36178, 86, M, and BBN_23398, 83, M) were deparaffinized in two changes of xylene of 5 minutes each, followed by rehydration through a graded series of ethanol (100%, 96%, 70%, 50%) for 5 minutes at each concentration and finally rinsed in distilled water for 5 minutes. Acid antigen retrieval was performed using citrate buffer (10 mM, pH 6.0, Tween 20, 0.05%) in a microwave oven. Tissue sections were submerged in the buffer and heated for 3 minutes at 95°C. After cooling to room temperature, the sections were washed in phosphate-buffered saline (PBS, pH 7.4) three times for 5 minutes.

Endogenous peroxidase activity was quenched by incubating the sections in 3% hydrogen peroxide (H₂O₂) in water for 10 minutes at room temperature, followed by two washes in PBS for 5 minutes each. To reduce nonspecific binding, sections were incubated with normal serum (from the species of the secondary antibody host) provided by the kit (ImmPRESS® HRP Goat Anti-Rabbit IgG Polymer Kit and ImmPRESS® HRP Horse Anti-Mouse IgG Polymer Kit, MP-7451 and MP-7402, Vector Laboratories) for 2 hours at room temperature.

Tissue sections were then incubated with the primary antibody (pE11, Synaptic Systems, 1:200 dilution or 4G8, Biolegend, 1:500 dilution) diluted in PBS with 1% bovine serum albumin (BSA) and 0.1% Triton X-100. Sections were incubated overnight at 4°C in a humidified chamber.

The day after, sections were rinsed in PBS three times for 5 minutes each. Sections were then incubated with the appropriate HRP conjugated secondary antibody belonging to the kit for 1 hour at room temperature. After washing with PBS (3 × 5 minutes), immunoreactivity was visualized using 3,3’-diaminobenzidine (DAB) chromogen solution prepared according to the manufacturer’s instructions (DAB Peroxidase Substrate, SK-4100, Vector laboratories). Sections were incubated with DAB for 4-5 minutes, monitoring the color development under a light microscope. The colorimetric reaction was stopped by rinsing the slides in distilled water.

For counterstaining, sections were immersed in hematoxylin for 3 minutes, rinsed in tap water, and dehydrated through graded alcohols (50%, 70%, 96%, and 100%) followed by xylene. Finally, sections were coverslipped with a permanent mounting medium (Permount mounting medium, SP15-100, Fisher Chemical).

Additionally, we performed in parallel the same procedure on a sample from a control individual as negative control, i.e. we treated the sections as described above, omitting only the addition of any primary antibody to the solution prior to the overnight incubation.

Immunostained sections were then imaged using a light microscope (Nikon Eclipse600) equipped with a DS-Fi3 color camera (Nikon, Tokyo, Japan) and a double LED light O-ring.

### PROTEOSTAT aggresome staining

To detect intracellular protein aggregation, we used the PROTEOSTAT® Aggresome detection kit (ENZ-51035, Enzo Life Sciences). In short, following the immunohistochemical detection of 4G8, slides were incubated for 30 minutes in a dark humidified box with Aggresome staining solution (1:500). Next, slides were washed three times in PBS, followed by mounting in prolong gold. Imaging was done using Ex/Em wavelengths of 550 – 565-600nm, respectively.

### Used antibodies

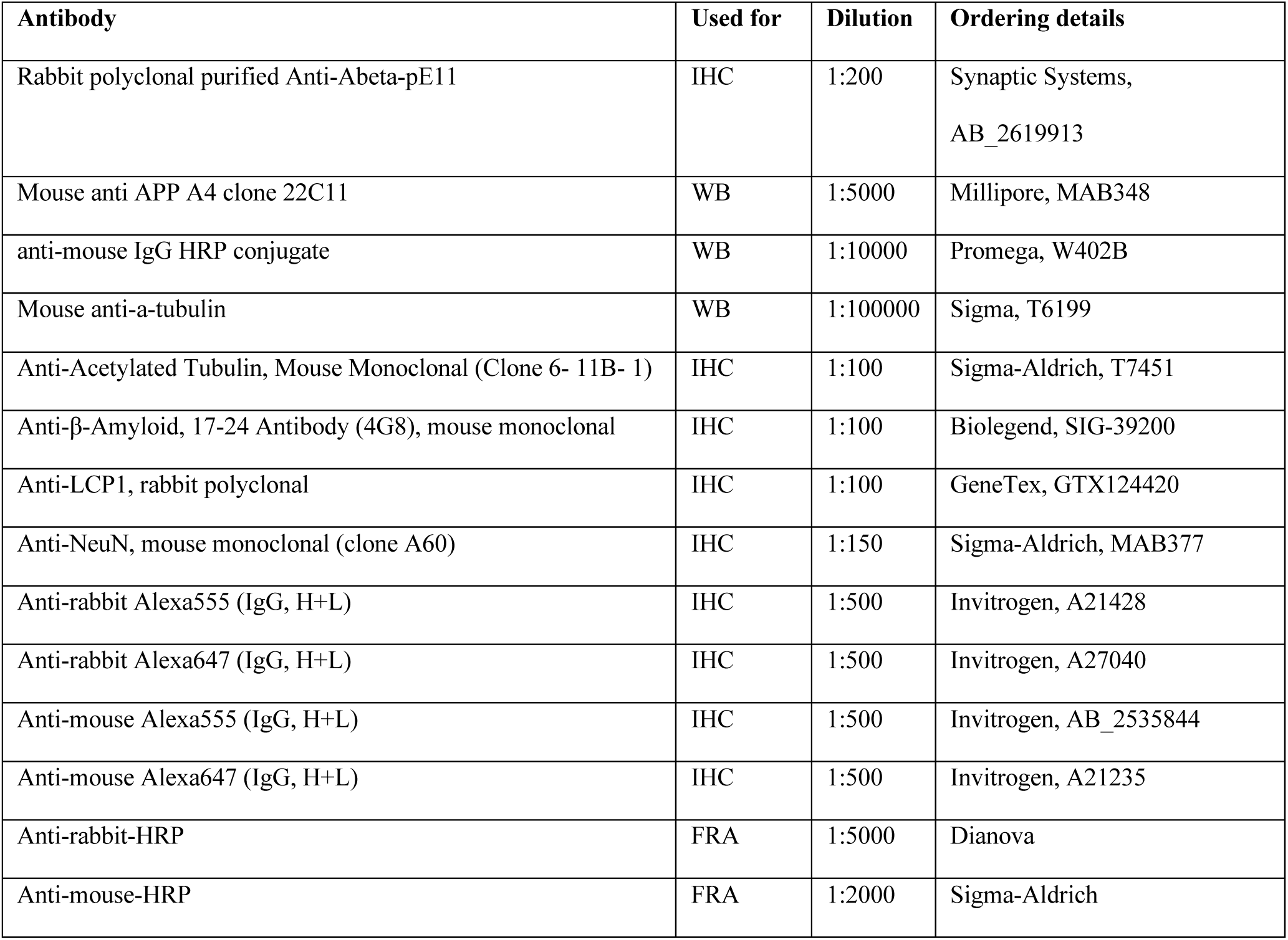

### RNA sequencing

Brain dissection was followed by RNA isolation using the TRIzol reagent (Life Technologies, Carlsbad, CA, USA) according to the manufacturer’s instructions. RNA integrity and concentration were measured using Agilent 4200 TapeStation System in combination with RNA ScreenTape (Agilent Technologies, Santa Clara, CA, US) ensuring high quality RNA (RNA Integrity Numbers > 8) for library preparation. Libraries for Illumina next-generation sequencing technology^86^ were prepared from 200 ng of total RNA using NEBNext Ultra II Directional RNA Library Preparation Kit in combination with NEBNext Poly(A) mRNA Magnetic Isolation Module and NEBNext Multiplex Oligos for Illumina (Unique Dual Index UMI Adaptors RNA) following the manufacturer’s instructions (New England Biolabs, Ipswich, MA, US). Library quantification and quality assessment were performed using Agilent 4200 TapeStation System with D5000 ScreenTape. Libraries were then pooled and sequenced on NovaSeq 6000 platform (Illumina, San Diego, CA, US) using the SP 100 cycle v1.5 kit with a 51-cycle paired-end read configuration. A minimum sequencing depth of 33.2 million reads per sample was achieved. Sequencing data were converted to FASTQ format using bcl2fastq (v2.20.0.422).

### RNA sequencing data analysis

Raw sequencing data (FASTQ files) were evaluated for quality using FastQC (v0.11.9). Adapter sequences, poly-A sequence and low-quality bases were removed using cutadapt (v4.2). High-quality reads were aligned to the reference genome (*Nothobranchius furzeri*, MPIA_NFZ_2.0) using STAR (2.7.6a) and PCR duplicates were removed based on unique molecular identifiers by umi-tools (1.1.1, parameter: --chimeric-pairs discard --unpaired-reads discard –paired). Gene-level quantification was performed using featureCounts (v2.0.3, parameter: -s 2 -p -B --countReadPairs) based on annotated gene features (obtained through personal communication with David Willemsen, corresponding to^87^. Differentially expressed genes were obtained using the R package EdgeR^88^ and gene ontologies were obtained using the online tool DAVID^89^.

### TUNEL staining

To visualize apoptotic cells in the killifish brain, cryosections (fixed in 4% PFA, sectioned 10um thick) were stained using the In Situ Cell Death Detection Kit, TMR rot (12156792910) from Roche. In short, cryosections were air dried for 15 seconds, and washed in PBS + 0,2% Triton-X100 for 3 times 10 minutes. Three washes in PBS, of each 5 minutes, were followed by placing the slides in a humidified box, and the brain sections of interest were encircled by a hydrophobic pen. Next, TUNEL staining solution was added on top of the sections which were placed, inside of the humidified box, in a 37°C incubator for 60 minutes. Afterwards, the slides were washed 3 times for 5 minutes in PBS and incubated with PBS+ DAPI (1:000 dilution) for 20 minutes in the humidified box. Last, the slides were washed 3 times for 5 minutes with PBS and mounted using Prolong Gold. Imaging and quantification were performed as described for immunohistochemistry, utilizing TUNELs excitation spectrum at 540nm and a emission spectrum between 570-620nm.

### Western blot analysis

Liquid frozen brains of killifish appa -/- and age matched controls of 6-months of age were lysed in 300µls 4xLämmli sample buffer (4 ml 20% SDS, 4 ml glycerol, 1ml ß-mercaptoethanol, 1 pinch bromophenol blue, 1,25 ml 1M Tris pH 7,6) using a tissue homogenizer. After boiling for 10 min at 95°C while shaking at 600 rpm, the samples were centrifuged 1 min at 11000 rpm. 20µls of the total lysate were loaded on Schägger gels. After 60 mins transfer to nitrocellulose blotting membrane (0,1 µm, Amersham^TM^ Protran, product 10600005) at 400 mA, the membrane was boiled for 5 mins in 1 x PBS, then incubated in 1 x PBS for 30 mins at room temperature before blocking with 0,2% Tropix I-Block (Applied Biosystems) for 1 hr at room temperature on a rotating platform. The incubation in primary antibody 22C11 (Mouse anti APP A4 clone 22C11, Millipore MAB348, 1:5000 in I-Block) was done overnight at 4°C. After removal of the primary antibody, the membrane was washed 4 x 15 mins in I-Block, followed by 1 hr incubation in a 1:10000 dilution of the secondary antibody (anti-mouse IgG HRP conjugate, W402B, Promega) at room temperature. The membrane was washed 4 times 15 mins and 3 times 10 mins with 1 x PBS before detection of bound antibodies with ECL plus (Pierce ECL Plus Western Blotting substrate, Thermo Scientific). After images were acquired using the LAS-4000 image reader (Fujifilm Life Science), the membrane was washed 4 x 15min in PBST, blocked in 3%milk for 1hr at room temperature and then incubated with the primary anti-a-tubulin antibody T6199 (Sigma) 1:100 000 in I-Block overnight at 4°C. After removal of the anti-a-tubulin antibody antibody, the membrane was washed 4 x 15 min in I-Block, followed by 1 hr incubation in a 1:10000 dilution of the secondary antibody (anti-mouse IgG HRP conjugate, W402B, Promega) at room temperature. The membrane was washed 4 x 15 min and 3 x 10 min with 1 x PBS before detection of bound antibodies with ECL plus (Pierce ECL Plus Western Blotting substrate, Thermo Scientific). Images of 22C11 and a-tubulin were quantified with the software ImageJ. For each group biological triplicates were quantified.

### Thioflavin-T assay

Whole brain protein lysates were produced using the same protocol as described to LC-MS. Black and transparent bottom 96-well plate (Greiner 96 F-bottom, Thermo Fisher #165305) were used, as well as the CLARIOstar Plus (BMG Labtech) plate reader. Thioflavin-T powder was bought from Sigma-Aldrich (T3516). To produce the Thioflavin-T (ThT) working solution, 80mg of ThT-powder was added to 25mL of dPBS (Gibco, [-] Cacl [-] MgCl) to make a 10mM stock. The solution was vortexed and incubated for 1 hour at room temperature. After the almost complete dissolvement of the ThT, 10mL of the ThT solution was filtered through a 0.2μm filter using a syringe. Next, 200μl of the filtered ThT solution was added to 1700μl of dPBS, to make the working solution of 50μM, and kept in the dark until further use. Next, protein lysates (49μl of 1.325μg/μl) was added to the 96-well plate (F-bottom). Using a multichannel pipet, 1μl of the ThT working solution (50μM) was added to the wells, and the plate was entered in the plate reader directly. The following imaging settings were used: Microplate: Greiner 96 F-bottom; Presets: CFP; Exitation: 430-20nm; Dichroic: auto 455nm; Emission: 480-20nm; Well scan: none; Optic: Bottom optic; Settling time: 0.2; Flying mode: off; No. of kinetic windows: 1; No. of cycles: 10; No. of flashes: 50; Cycle time: 60; Shake: Before first cycle; double orbital, 100rpm, 10sec. Values were recorded after 3 minutes and 30 seconds. Biological replicates were used in duplo, and averaged values were plotted.

### Filter retardation assay

Brain tissues were transferred to 2 mL Precellys tubes containing a mix of 1.4 mm and 2.8 mm ceramic (zirconium oxide) beads, in 200 µL of BPS-PIS buffer. Two cycles of mechanical homogenization were carried out at 6000 rpm for 30 seconds each, with a 30-second break on ice between each cycle, using the Precellys 24 lysis and homogenizer (Bertin, France). Subsequently, the tubes were centrifuged at 3000 rpm for 5 min at 4°C. The resulting supernatant was carefully transferred to a new 1.5 mL low-binding tube. An equal volume of 2X lysis buffer was added to the homogenate and sonicated (Bioruptor Plus, Diagenode, Belgium) for 20 cycles (60 sec ON/30 sec OFF) at 4°C. The protein concentration in the lysate was quantified using the Pierce™ BCA Protein Assay Kit (Thermo Fisher Scientific, Cat. no. 23227). For the filter retardation analysis, we adapted a previously published protocol^57^. In brief, 100 µg of protein was loaded in duplicate onto a 0.2 µm cellulose acetate membrane (Cytiva, Cat. no. 10404180) that had been pre-equilibrated in 1X TBS. The membrane was washed twice with a wash buffer (150 mM NaCl and 10 mM Tris, pH 8.0), then blocked with 5% milk. This was followed by incubation with primary antibodies: Rabbit-anti-pE11 (1:500, Synaptic Systems) or Mouse-anti-4G8 (1:1000, BioLegend), overnight. Subsequent incubation with secondary antibodies was performed using anti-rabbit-HRP (1:5000, Dianova) and anti-mouse-HRP (1:2000, Sigma-Aldrich) for 4 h. The images were captured in Amersham imager 600, and quantification was performed using Fiji ImageJ software. Buffers: BPS-PIS: 250 µL PBS; 5 µL Protease inhibitor cocktail; 25 µL PhosSTOP. Lysis Buffer (2X): 225 µL PBS; 25 µL SDS (10%); 12.5 µL HEPES (2M). Tris-buffered saline (TBS) 10X, pH 8: 200 mM Tris base; 1.5 M NaCl. Tris-buffered saline with Tween (TBS-T): 1X TBS; 0.1% Tween 20.

### CRISPR/Cas9 transgenesis

CRISPR/Cas9 tool was used to edit the genome of the turquoise killifish. Guide RNA templates were designed using an online program CHOPCHOP (https://chopchop.cbu.uib.no/). Exon 2 of *appa* was targeted using two different gRNAs, and Exon 3 of *appb* was targeted using three different gRNAs. Templates of ssDNA target-specific oligonucleotides were used for sgRNA synthesis. Used guide RNA’s (regions in bold indicate the complement sequence of the target):

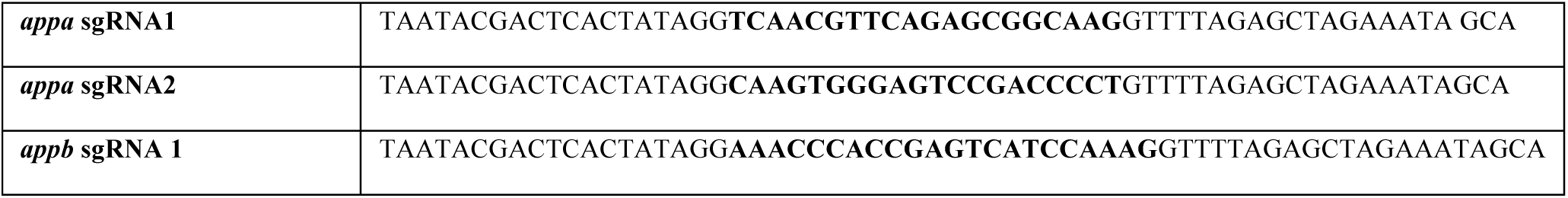
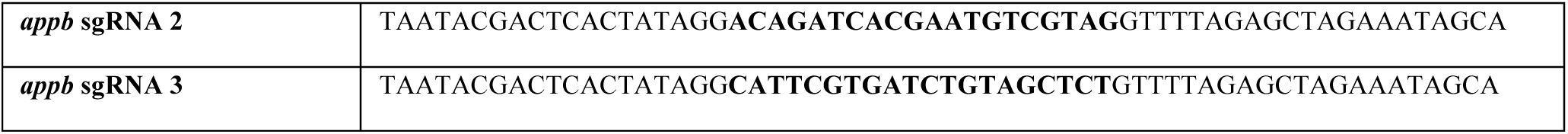

The template ssDNA target-specific oligonucleotide to synthesise gRNA was designed by adding an overlapping nucleotide sequence to the N-terminus and T7 promoter sequence to the C-terminus. The PAM sequence (NGG) was removed from the sequence. Templates were synthesised and delivered by the Integrated DNA Technologies (https://eu.idtdna.com/pages/products/custom-dna-rna/dna-oligos). The sgRNAs were synthetised using the EnGen® sgRNA Synthesis Kit (New England BioLabs, USA, #E3322V), according to their standardized protocol. The synthesised sgRNAs was cleaned up with the Monarch® RNA Cleanup Kit (50μg, New England BioLabs, USA, #T2040L), following the instructions. For CRISPR/Cas9 injections, concentration of sgRNA solution was adjusted to 300 ng/μL. The CRISPR/Cas9 and gRNA reaction mix was prepared with EnGen® Spy Cas9 NLS (20 μM, New England Biolags, USA), by mixing 1 μL of Cas9 NLS, 1 μL of each purified sgRNA (300 ng/μL), 2 μL Phenol Red for visualisation and nuclease free water up to 10 μL (kept on ice). The needles were pulled from glass capillaries 1.0 OD x 0.58 ID x 100 L mm using a micropipette puller machine with the program previously established in the lab. One- or two-cell stage embryos were placed and fixed in the agarose plate and injected with a small amount of reaction mix directly into the embryos one at a time. The embryos were transferred to the clean methylene blue solution, checked daily and further handled as previously described for embryo maintenance. To genotype CRISPR/Cas9 injected fish and their progeny, first fish DNA had to isolated from 2-4 scales which were collected from the posterior part of the body of sexually matured fish (minimum 5-6 weeks of age). The scales were transferred to 50 μL of SEL buffer (1M KCl, 1M MgCl_2_, 1M Tris pH 8,3, 10% Igepal, 10% Tween-20, 0,1% Gelatine in miliQ H_2_O) and 1 μL of Proteinase K. The scales were incubated in the SEL isolation buffer and Proteinase K at 60°C for 1 h, 90°C for 15 min and 12°C until tubes were taken out of the machine. The isolated DNA was used as a template for PCR amplification of the target region of interest. *Taq* 2X Master Mix was used for the PCR reaction. PCR details. Forward primer: CCTCTCTTTCAGGTCCCCAC. Reverse primer:

TTTTCTGCTCTGCT TTTAGGC. Amplicon Length: 209 bp. CONDITIONS: **(1x)** 3:00 95.0C **(35x)** 0:45 95.0C 0:30 55.0C 0:45 72.0C (**1x)** 5:00 68.0 ∞ 10.0 PCR amplicons were purified using Sera-Mag SpeedBead Carboxylate-Modified Magnetic Particles (Hydrophylic) magnetic beads. Purified DNA was sent for Sanger sequencing to the Eurofins Genomics (https://eurofinsgenomics.eu/).

### Sample preparation for proteomics analysis

Before lysis, lysis buffer (fc 4% SDS, 100 mM HEPES, pH 8.5, 50 mM DTT) was added to each samples. Samples were then boiled at 95°C for 7 min and sonicated using a tweeter. Reduction was followed by alkylation with 200 mM iodoacetamide (IAA, final concentration 15 mM) for 30 min at room temperature in the dark. Samples were acidified with phosphoric acid (final concentration 2.5%), and seven times the sample volume of S-trap binding buffer was added (100 mM TEAB, 90% methanol). Samples were bound on 96-well S-trap micro plate (Protifi) and washed three times with binding buffer. Trypsin in 50 mM TEAB pH 8.5 was added to the samples (1 µg per sample) and incubated for 1 h at 47°C. The samples were eluted in three steps with 50 mM TEAB pH 8.5, elution buffer 1 (0.2% formic acid in water) and elution buffer 2 (50% acetonitrile and 0.2% formic acid). The eluates were dried using a speed vacuum centrifuge (Eppendorf Concentrator Plus, Eppendorf AG, Germany) and stored at −20° C. The samples were loaded on Evotips (Evosep) according to the manufacturer’s instructions. In short, Evotips were first washed with Evosep buffer B (acetonitrile, 0.1% formic acid), conditioned with 100% isopropanol and equilibrated with Evosep buffer A. Afterwards, the samples were loaded on the Evotips and washed with Evosep buffer A. The loaded Evotips were topped up with buffer A and stored until the measurement.

### LC-MS Data independent analysis (DIA)

Peptides were separated using the Evosep One system (Evosep, Odense, Denmark) equipped either with a 15 cm x 150 μm i.d. packed with a 3 μm Reprosil-Pur C18 bead column (Evosep Endurance, EV-1109, PepSep, Marslev, Denmark) or with a a 15 cm x 150 μm i.d. packed with a 1.5 μm Reprosil-Pur C18 bead column (Evosep Performance, EV-1137, PepSep, Marslev, Denmark) heated at 45°C with a butterfly sleeve oven (Phoenix S&T, Philadelphia, USA). The samples were run with a pre-programmed proprietary Evosep gradient of 44 min (30 samples per day) using water and 0.1% formic acid and solvent B acetonitrile and 0.1% formic acid as solvents. The LC was coupled to an Orbitrap Exploris 480 (Thermo Fisher Scientific, Bremen, Germany) using PepSep Sprayers and a Proxeon nanospray source. The peptides were introduced into the mass spectrometer via a PepSep Emitter 360-μm outer diameter × 20-μm inner diameter, heated at 300°C, and a spray voltage of 2 kV was applied. The injection capillary temperature was set at 300°C. The radio frequency ion funnel was set to 30%. For DIA data acquisition with endurance column, full scan mass spectrometry (MS) spectra with a mass range of 350–1650 m/z were acquired in profile mode in the Orbitrap with a resolution of 120,000 FWHM. The default charge state was set to 2+, and the filling time was set at a maximum of 60 ms with a limitation of 3 × 106 ions. DIA scans were acquired with 40 mass window segments of differing widths across the MS1 mass range. Higher collisional dissociation fragmentation (normalized collision energy 29%) was applied, and MS/MS spectra were acquired with a resolution of 30,000 FWHM with a fixed first mass of 200 m/z after accumulation of 1 × 106 ions or after filling time of 45 ms (whichever occurred first). Data were acquired in profile mode. For DIA data acquisition for Performance column, full scan mass spectrometry (MS) spectra with a mass range of 350– 1650 m/z were acquired in profile mode in the Orbitrap with a resolution of 120,000 FWHM. The default charge state was set to 2+, and the filling time was set at a maximum of 20 ms with a limitation of 3 × 106 ions. DIA scans were acquired with 45 mass window segments of differing widths across the MS1 mass range. Higher collisional dissociation fragmentation (normalized collision energy 30%) was applied, and MS/MS spectra were acquired with a resolution of 30,000 FWHM with a fixed first mass of 200 m/z after accumulation of 1 × 106 ions or after filling time of 45 ms (whichever occurred first). For data acquisition and processing of the raw data, Xcalibur 4.5 (Thermo) and Tune version 4.0 were used.

### Proteomic data processing

DIA raw data were analysed using the directDIA pipeline in Spectronaut (v.17 and v.18, Biognosysis, Switzerland). The data were searched against an in house species-specific database (*N.furzeri*, 59154 entries, Nfu_20150522, annotation nfurzeri_genebuild_v1.150922) with a list of common contaminants appended (247 entries). The data were searched with the following variable modifications: oxidation (M) and acetyl (protein N-term). A maximum of 2 missed cleavages for trypsin and 5 variable modifications were allowed. The identifications were filtered to satisfy an FDR of 1% at the peptide and protein levels. Relative quantification was performed in Spectronaut for each paired comparison using the replicate samples from each condition. Precursors were percentile filtered at 0.2%. The data (candidate table) and data reports (protein quantities) were then exported, and further data analyses and visualization were performed with R studio using in-house pipelines and scripts. Differentially abundant proteins were defined as having a q-value <0.

### Active avoidance test

We assessed the decline in learning ability of treated fish using an active avoidance behavioral test which was developed based on a test previously employed in killifish ^39,41^. The setup consisted of two circular compartments connected through a small opening. Fish were acclimatized for 15 minutes in water from their home tank before testing began. A red light served as the conditioned stimulus, followed by air bubbles as the negative stimulus. Fish learned to avoid the negative stimulus by crossing the small opening into the opposite compartment when the red light appeared. The red light was presented for 30 seconds, with bubbles introduced after 15 seconds if the fish had not moved. Trials were counted as ‘success’ if the fish crossed the hurdle within 15 seconds of the red-light onset (before the introduction of bubbles). The cycle of stimulus, bubbles, and rest repeated 40 times per fish. Each fish underwent the test individually and was sacrificed immediately afterward. The entire test lasted one hour per animal. The water temperature was kept stable between 26.5°C and 28.0°C using a standard aquarium heater, which was important as varying temperatures could impact the performance of the fish.

### Hybridizing Chain Reaction

Probe pair pools targeting *fos* and *neun* transcripts were generated and validated as previously described^35,90^. The probe sequences can be found in **Table S5**. The selected probe pairs were ordered from Integrated DNA Technologies, Inc. (IDT). The HCR protocol (HCRv3.0) was followed as previously published^91^ with adaptations for cryosections as described by Van Houcke et al. (2021)^24^. Cryosections of 10um thick were used. The amplification of the signal was done for *fos* transcripts with HCR Amplifier B1, Alexa Fluor 647 and *neun* transcripts with HCR Amplifier B2, Alexa Fluor 546. Imaging was done with the Zeiss Axio Scan.Z1 microscope with a Plan-Apochromat 20x/0.8 M27 objective, Axiocam 506m imaging device and 1x Camera Adapter.

## Acknowledgements

We are thankful to the Core Facility Proteomics, Imaging, Histology & Electron Microscopy, and Next Generation Sequencing at the FLI. We are thankful to the Animal fish Facility of the FLI, with a special mention of Johannes Wilfert, as well as Matthias Werres and Christina Paetzold from the fish facility of the MPI-AGE. We would like to thank the group of Lutgarde Arckens at K.U. Leuven (Belgium), and especially Caroline Zandecki, for the help with setting up the hybridizing chain reaction experiment. We are thankful to the group of Randal Halfmann (Stowers Institute) for the expert advice and help with the interpretation of the data, as well as Mario Baumgart for his help with importing the *appa* mutant line to the FLI. D.E.M. de Bakker was financed by a Rubicon scholarship (452021116) from the Dutch Research Council (NWO) as well as by a research grant from the “Balance of the Microverse” Excellence Cluster from the Friedrich Schiller University in Jena. We acknowledge the contribution of the London Neurogenerative Diseases Brain Bank (which is part of the Brains for Dementia Research network, jointly funded by Alzheimer’s Research UK and the Alzheimer’s Society) that provided us with the human-derived brain samples utilized in this study. This project was funded by the MPI-AGE and from the FLI in Jena. We would like to thank all the past and current members of the Valenzano lab at the MPI-AGE and at the FLI for their continuous input and support.

## Data availability

The produced transcriptomic datasets will be made publicly available at the Gene Expression Omnibus from NCBI. To review GEO accession GSE281018, go to https://www.ncbi.nlm.nih.gov/geo/query/acc.cgi?acc=GSE281018. Enter token *cxybacccdzuzluz* into the box. Proteomics data will be made publicly available at MassIVE, which can be found following the links below. Proteomics data for Young (7 weeks) versus Old (6 months) *N. furzeri* brains: http://massive.ucsd.edu/ProteoSAFe/status.jsp?task=650a5a2e981540359f34b145717d2f7b Proteomics data for *N. furzeri appa* -/- versus *appa* +/+ brains: http://massive.ucsd.edu/ProteoSAFe/status.jsp?task=e7c08aa99a7a427c8ec6eb2623a9e5ec

## Author contributions

**DdB** planned the project, managed, designed, and executed most of the experiments including histology, behavioral analysis, and statistical and bioinformatical analysis, produced the figures, acquired funding, and wrote the manuscript. **MM** generation the appa KO line and histological analysis. Produced supplementary figures (S4-8). **KG** and **YR** performed the Filter Retardation Assay. **SB** performed histology on human samples. **FvB:** performed 22C11 Western Blot and critical contributed to the experimental design and interpretation. **BS** Critically contributed to the experimental design and interpretation. **LA** Assisted **DdB** with brain isolation and immunohistochemistry. **FSS** set up and performed HCR experiments. **OO** ran the AlphaFold model. **MB** supervised the transcriptomic data production and analysis. **EC** supervised the proteomic data production and analysis. **AA** shared resources and provided input on the manuscript. **IM** contributed to the manuscript draft. **AS** helped with project conceptualization and provided the pE11 antibody. **AC** aided with sample acquisition and critically contributed to the manuscript draft. **JK** supervised the biochemical analysis of protein aggregation and critically contributed to the manuscript draft. **DRV** Planned and supervised the project, developed the Bayesian models, wrote the manuscript, and acquired the funding.

---

**Figure S1.**
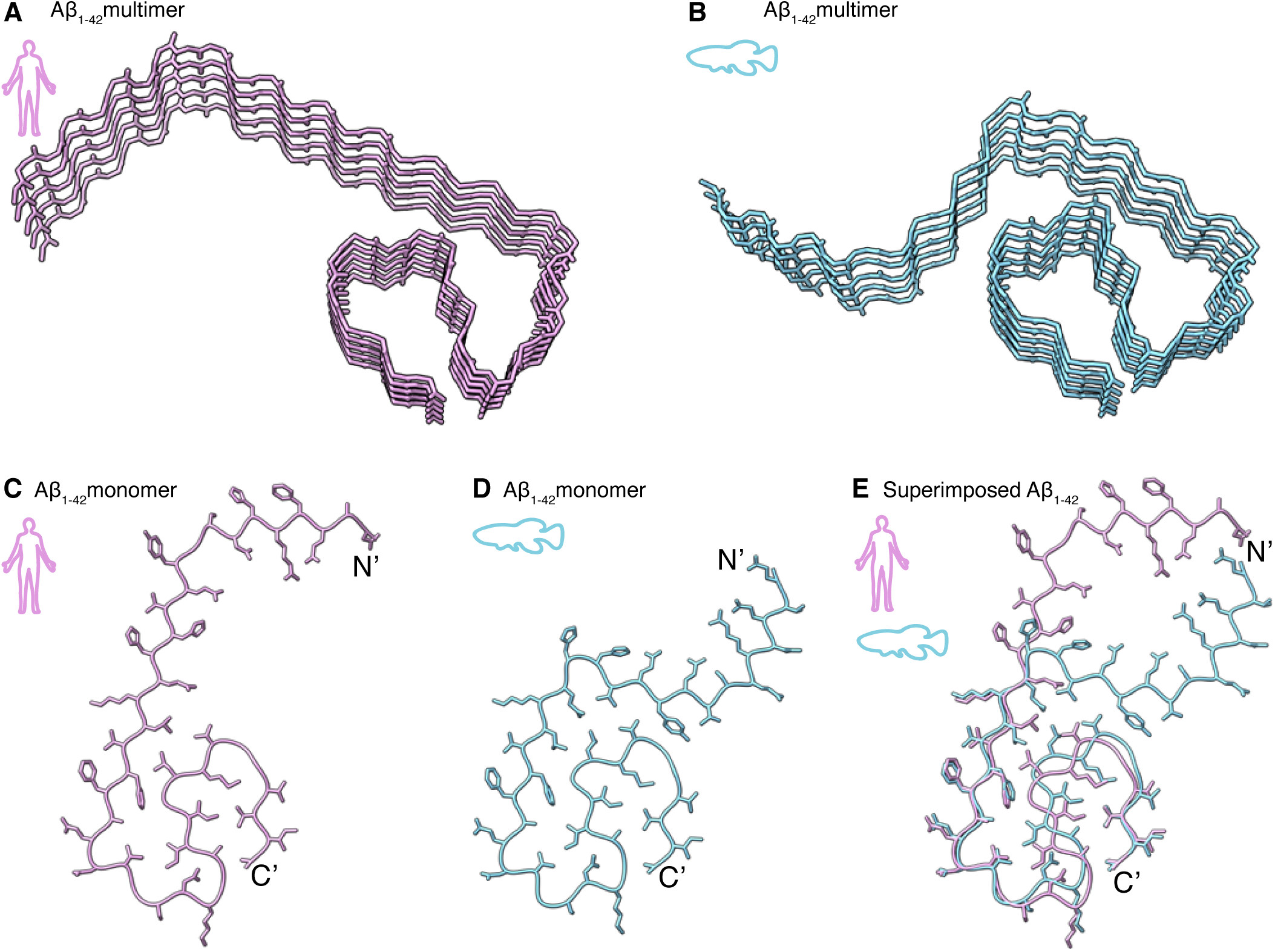
3D protein structure of amyloid beta (1-42) is conserved between turquoise killifish and humans. (**A**) Alphafold (V3) predicted stucture of human amyloid beta (1-42) in an aggregated state. (**B**) Alphafold-predicted stucture of killifish amyloid beta (1-42) derived from *appa* in an aggregated state. (**C**) Alphafold prediction of human amyloid beta monomer 1-42), as part of the multimer shown in (A). (**D**) Alphafold prediction of turquoise killifish amyloid beta monomer (1-42) derived from *appa*, as part of the multimer shown in (B). (**E**) Superimposed of the Alphafold-predicted multimer structures (only one monomer displayed) of human (pink) and killifish (blue) amyloid beta (1-42) (backbone r.m.s.d. for residues Q15-A42: 0.859 Angstrom).

**Figure S2.**
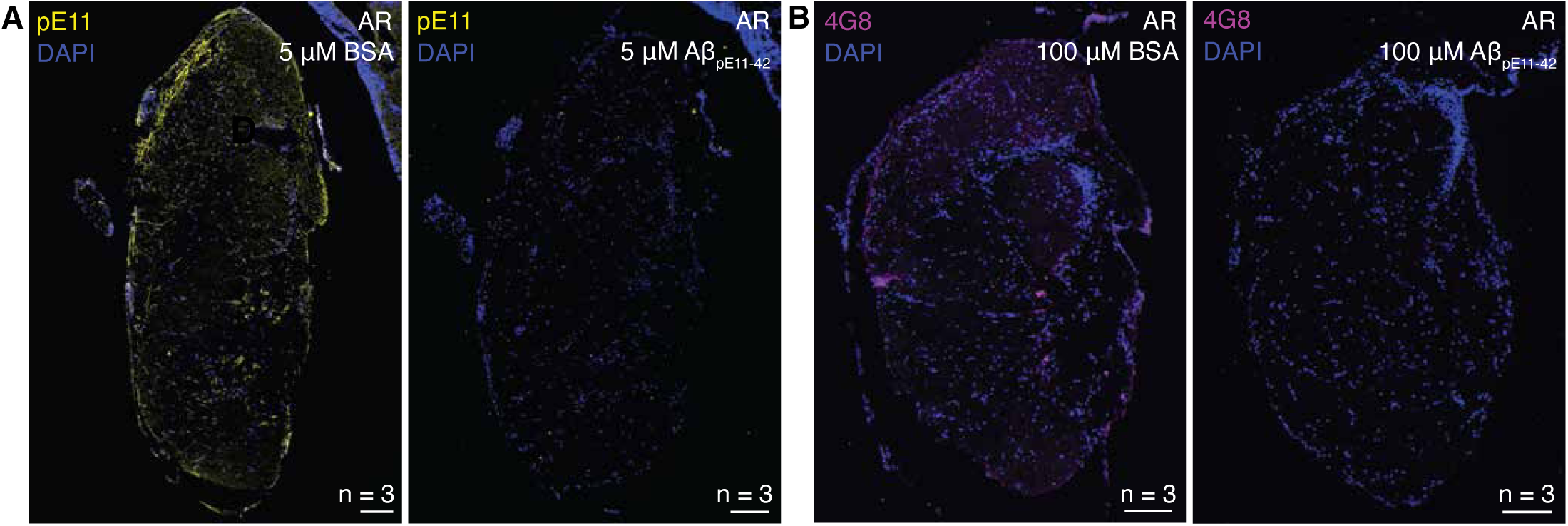
pE11 and 4G8 antibodies bind their targets in an epitope dependent manner. (A-B) Immunohistochemistry staining using the pE11 (A) or 4G8 (B) antibody, after a 30 minute incubation step of the primary antibody together with either BSA (left) or AβpE11_-42_ peptide (right). The anterior rhombencephalon (AR) was stained from three 6-month-old turquoise killifish per condition. Scalebars represent 100 µm.

**Figure S3.**
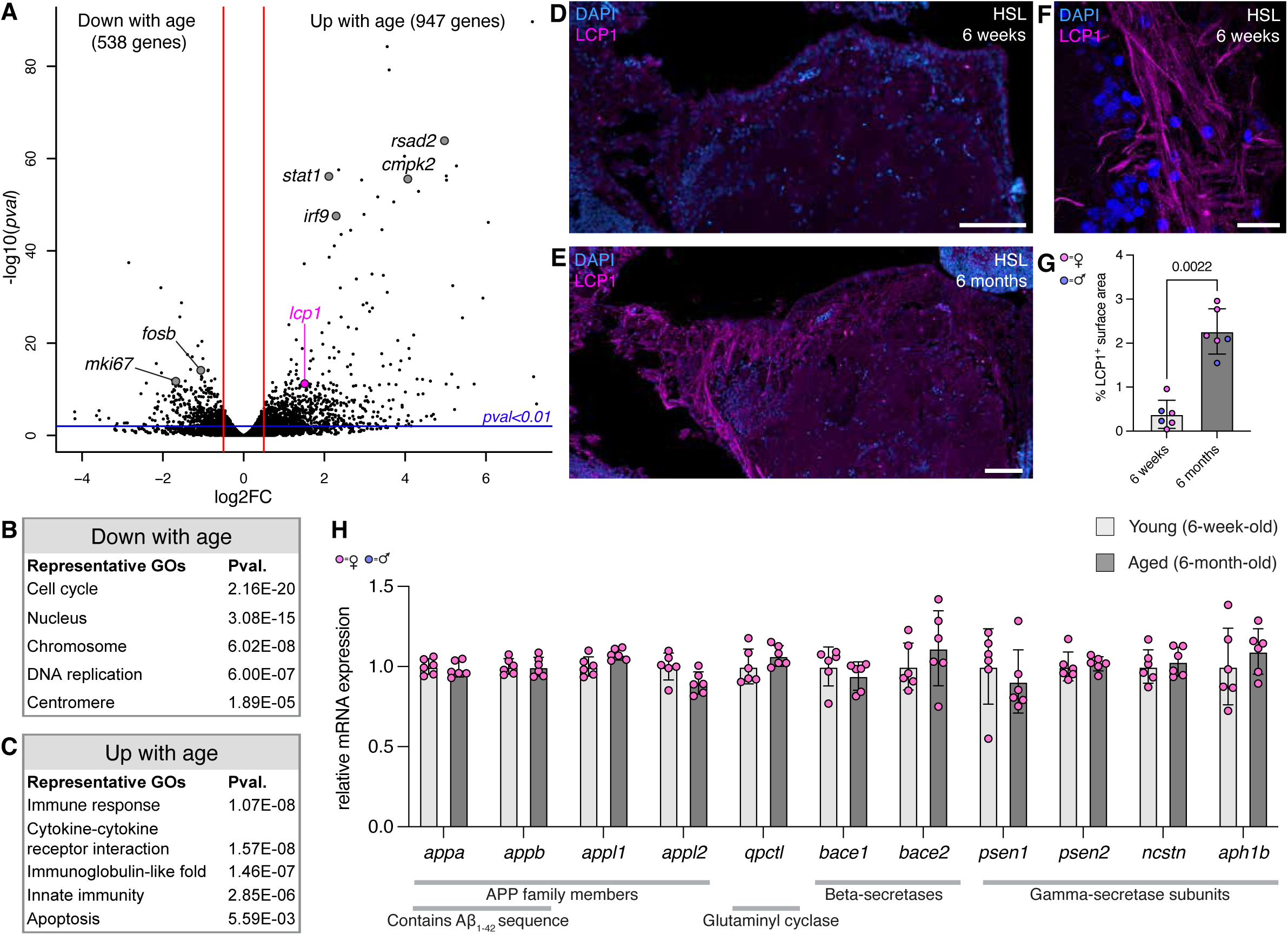
The turquoise killifish brain develops an age-related inflammatory state while expressing key players of Aβ biogenesis. (**A**) Volcano plot displaying differences in gene expression levels between young (6-week-old) and aged (6-month-old) female killifish brains. Differentially expressed genes are highlighted, representing relevant biological processes. Vertical red lines indicate log2FC of (-)0.5. (**B, C**) Representative gene ontologies produced from the differentially expressed genes between young and old brains (Log2FC > 0.5 / < −0.5 and pval < 0.01). (**D, E**) Immunohistochemistry staining targeting leukocyte marker Lymphocyte Cytosolic Protein 1 (LCP1) in sections of the hypothalamus superior lobe (HSL) from a young (D) and aged (E) individual. Scale bars represent 100 µm. (**F**) High-resolution image of the HSL of a 6-month-old killifish brain, showing the striated pattern of the LCP1 staining, which reflects binding to the actin cytoskeleton. Scale bar represents 20 µm. (**G**) Quantification of the percentage of LCP1+ covered surface area in the hypothalamus of young and aged brains. Significance was calculated using a Mann-Whitney test. Error bars represent standard deviation. (**H**) Expression levels of key players in Aβ biogenesis normalized to total mRNA counts per brain, plotted as relative values to the mean value of young (6-week-old) individuals. APP = Amyloid beta Precursor Protein. Error bars represent standard deviation.

**Figure S4.**
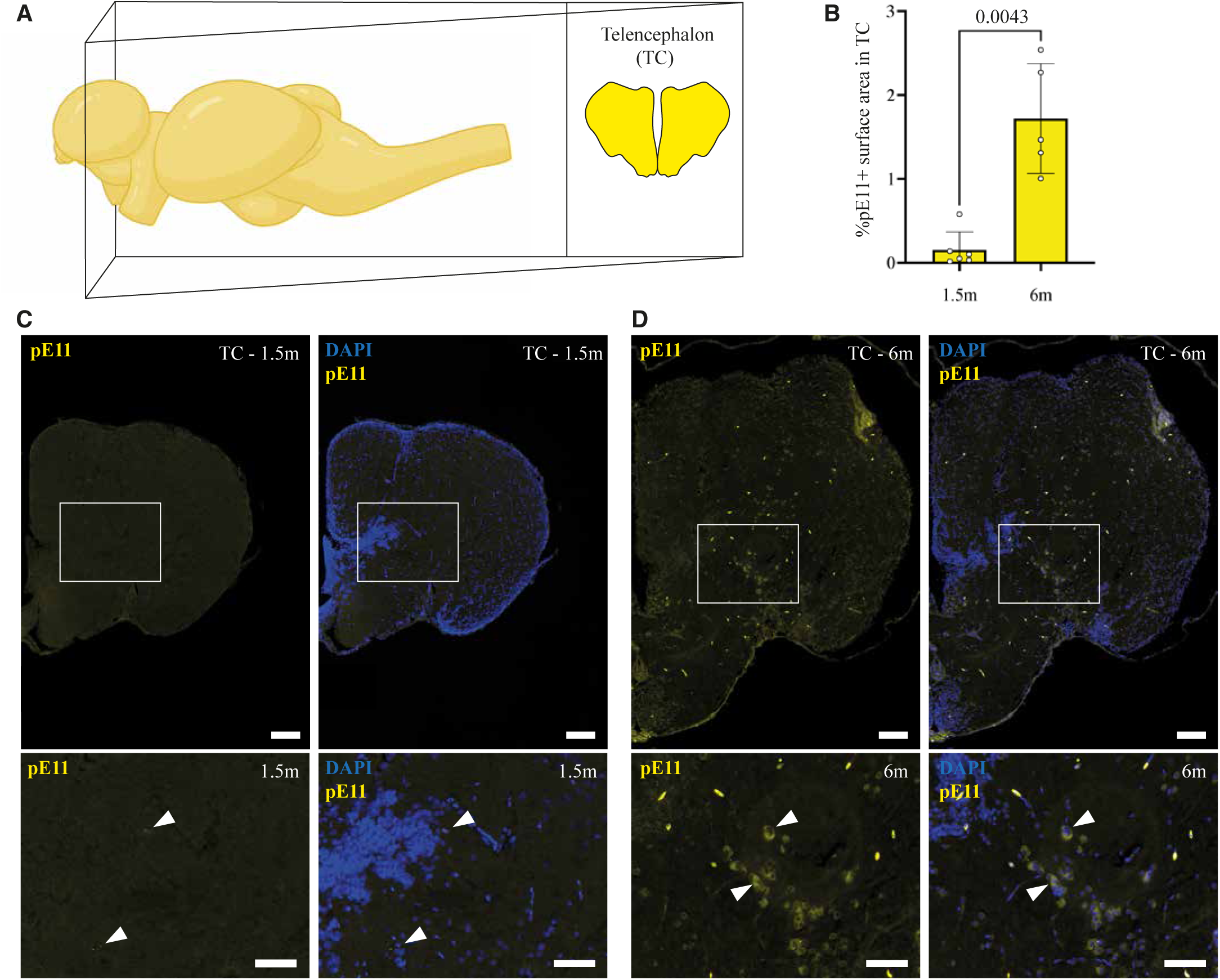
pE11 signal accumulates in the telencephalon in an age-dependent manner. (**A**) Schematic representation of the brain region quantified and shown in B-D. (**B**) Quantification of immunohistochemistry staining using the pE11 antibody on the telencephalon (TC) of young (6-week-old) and aged (6-month-old) killifish. Significance was calculated using a Mann-Whitney test. (**C, D**) Immunohistochemistry staining with the pE11 antibody (yellow) and DAPI (blue) in a 6-week-old (C) and 6-month-old (D) telencephalon. Arrowheads indicate examples of regions with perinuclear immunoreactivity. Scale bar represents 100 µm in the overview and 50 µm in the zoom-in.

**Figure S5.**
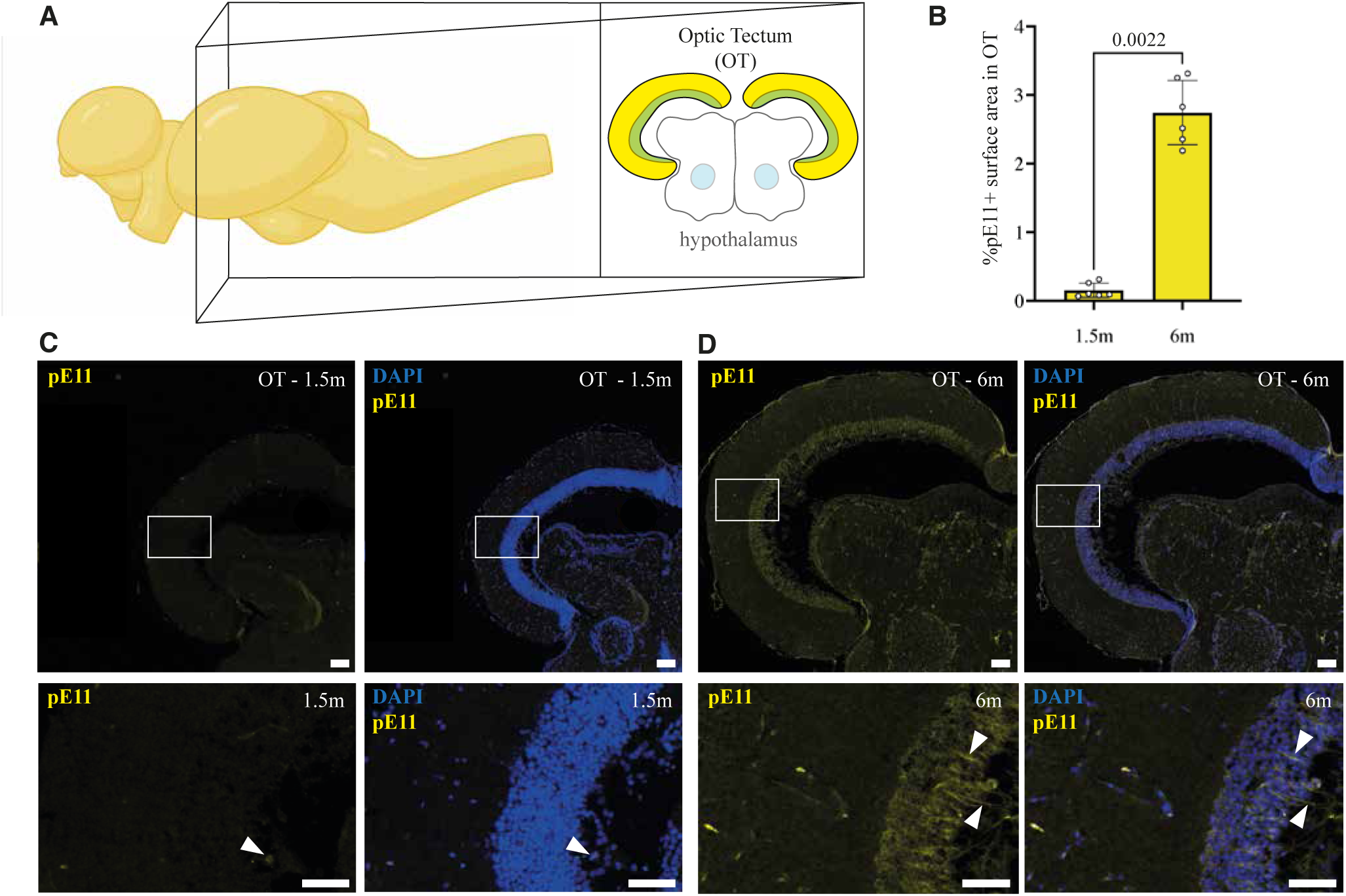
pE11 signal accumulates in the optic tectum in an age-dependent manner. (**A**) Schematic representation of the brain region quantified and shown in B-D. (**B**) Quantification of immunohistochemistry staining using the pE11 antibody on the optic tectum (TC) of young (6-week-old) and aged (6-month-old) killifish. Significance was calculated using a Mann-Whitney test. (**C, D**) Immunohistochemistry staining with the pE11 antibody (yellow) and DAPI (blue) in a 6-week-old (C) and 6-month-old (D) optic tectum. Arrowheads indicate regions with immunoreactivity. Scale bars represent 100 µm in the overviews and 50 µm in the zoom-ins.

**Figure S6.**
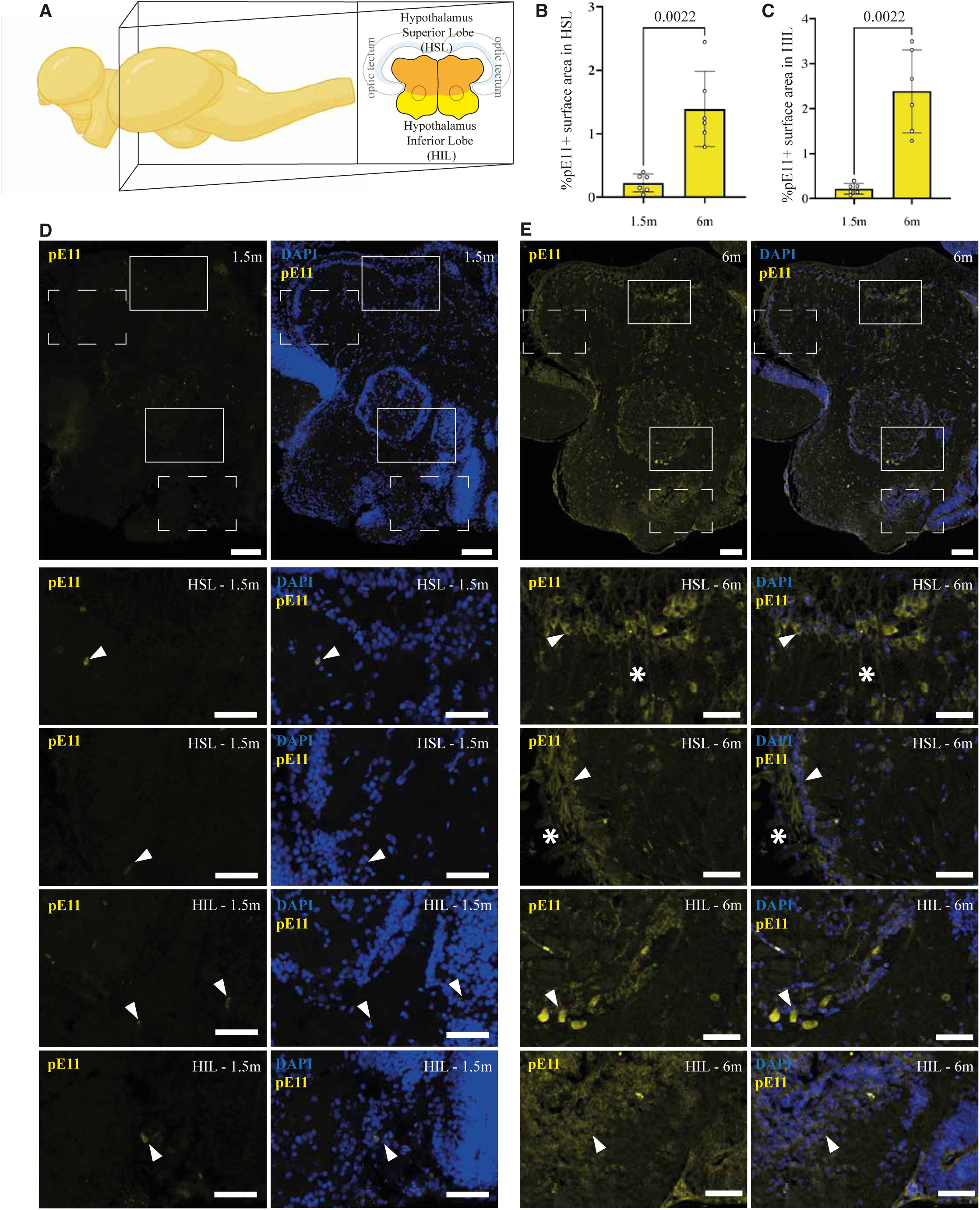
pE11 signal accumulates in the hypothalamus in an age-dependent manner. (**A**) Schematic representation of the brain regions quantified and shown in B-E. (**B,C**) Quantification of immunohistochemistry staining using the pE11 antibody on the hypothalamus superior lobe (HSL) and hypothalamus inferior lobe (HIL) of young (6-week-old) and aged (6-month-old) killifish. Significances were calculated using Mann-Whitney tests. (**D, E**) Immunohistochemistry staining with the pE11 antibody (yellow) and DAPI (blue) in 6-week-old (C) and 6-month-old (D) hypothalamus. Arrowheads indicate regions with perinuclear immunoreactivity. Asterisks indicate regions with filamentous immunoreactivity. Scale bars represent 100 µm in the overviews and 50 µm in the zoom-ins.

**Figure S7.**
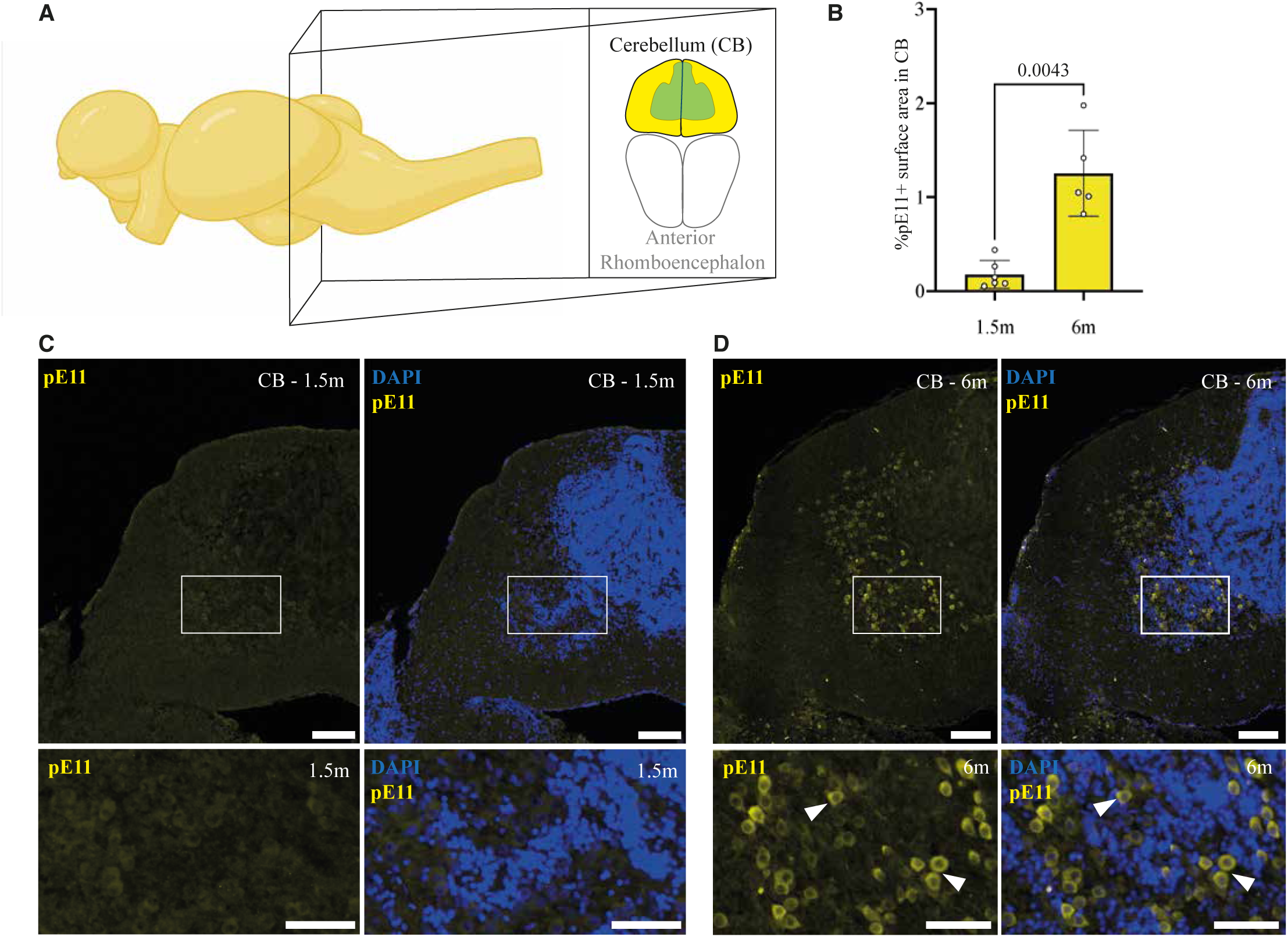
pE11 signal accumulates in the cerebellum in an age-dependent manner. (**A**) Schematic representation of the brain region quantified and shown in B-D. (**B**) Quantification of immunohistochemistry staining using the pE11 antibody on the cerebellum (CB) of young (6-week-old) and aged (6-month-old) killifish. Significance was calculated using a Mann-Whitney test. (**C, D**) Immunohistochemistry staining with the pE11 antibody (yellow) and DAPI (blue) in a 6-week-old (C) and 6-month-old (D) cerebellum. Arrowheads indicate examples of regions with perinuclear immunoreactivity. Scale bars represent 100 µm in the overviews and 50 µm in the zoom-ins.

**Figure S8.**
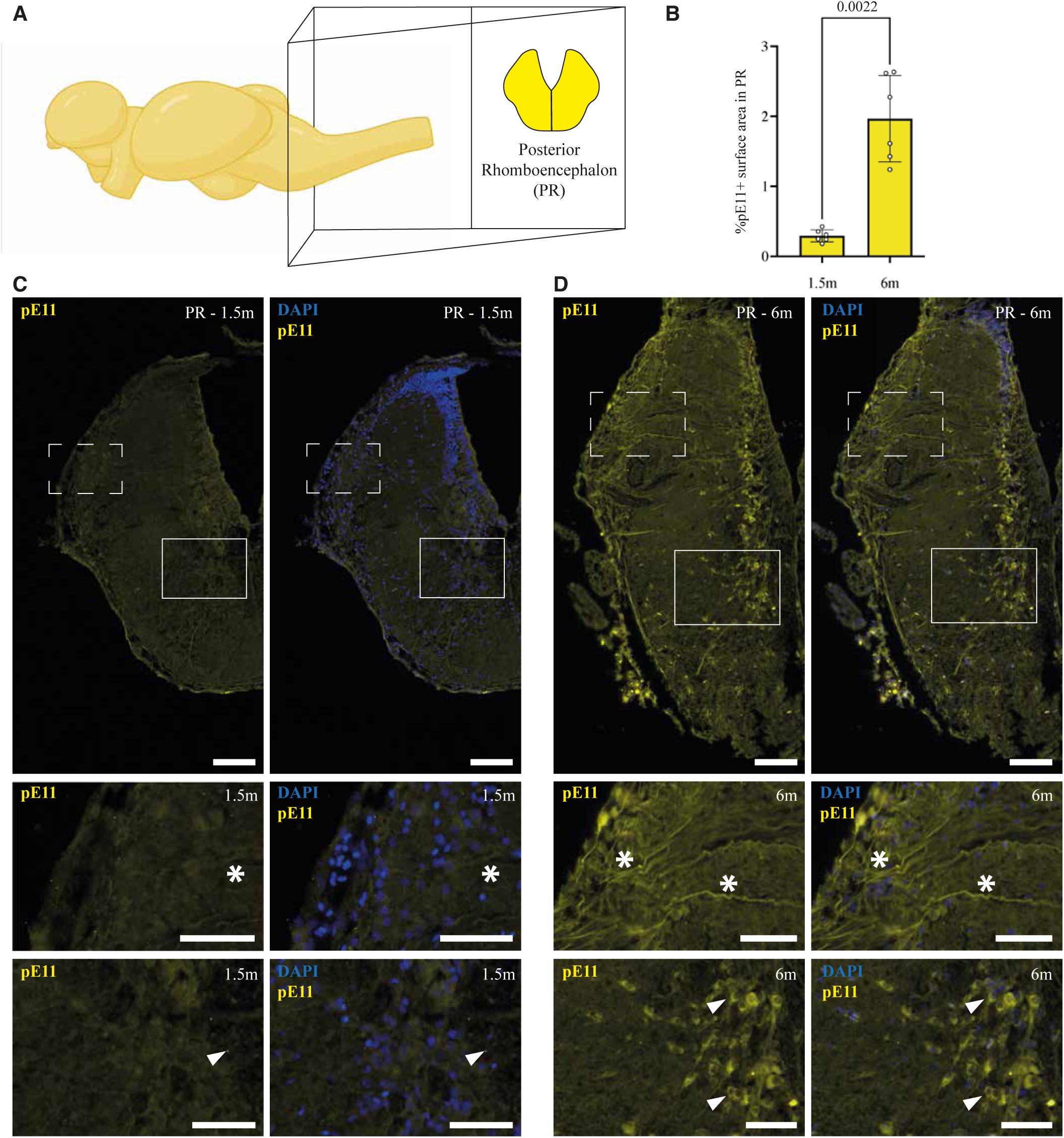
pE11 signal accumulates in the posterior rhomboencephalon in an age-dependent manner. (**A**) Schematic representation of the brain region quantified and shown in B-D. (**B**) Quantification of immunohistochemistry staining using the pE11 antibody on the posterior rhomboencephalon (CB) of young (6-week-old) and aged (6-month-old) killifish. Significance was calculated using a Mann-Whitney test. (**C, D**) Immunohistochemistry staining with the pE11 antibody (yellow) and DAPI (blue) in a 6-week-old (C) and 6-month-old (D) posterior rhomboencephalon. Arrowheads indicate examples of regions with perinuclear immunoreactivity. Asterisks indicate regions with filamentous immunoreactivity. Scale bars represent 100 µm in the overviews and 50 µm in the zoom-ins.

**Figure S9.**
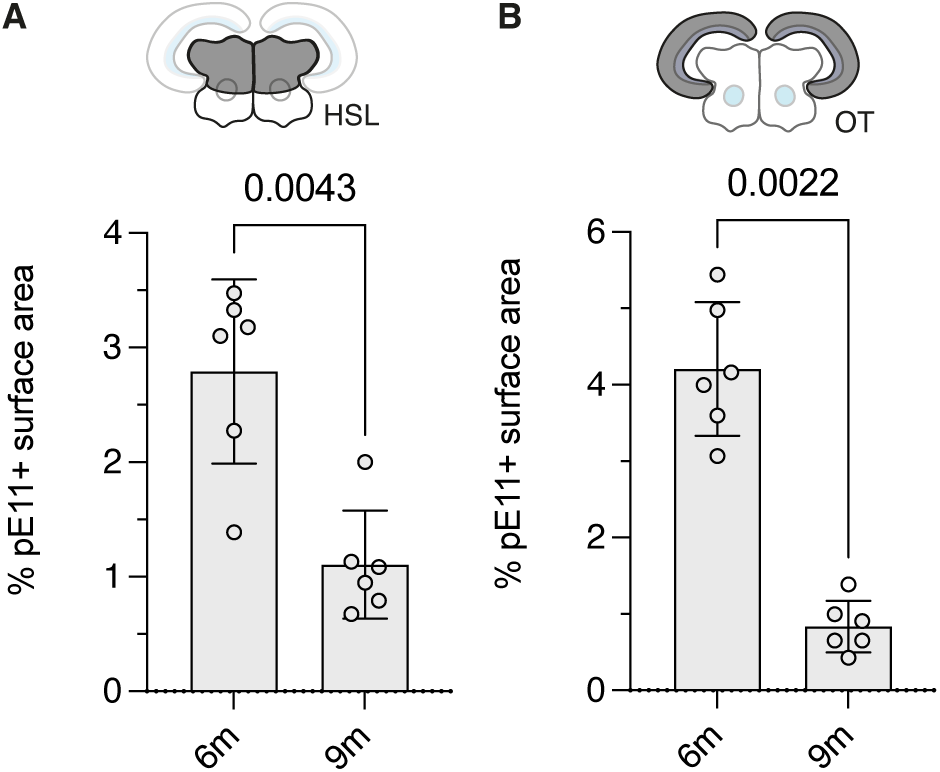
pE11 signal decreases in the turquoise killifish brain between 6- and 9-months of age. (**A, B**) The percentage surface area covered was quantified in an independent cohort of turqoise killifish (strain ZMZ1001) in the (A) hypothalamus superior lobe (HSL) and (B) the optic tetum (OT). Significance was calculated with a Mann-Whitney test.

**Figure S10.**
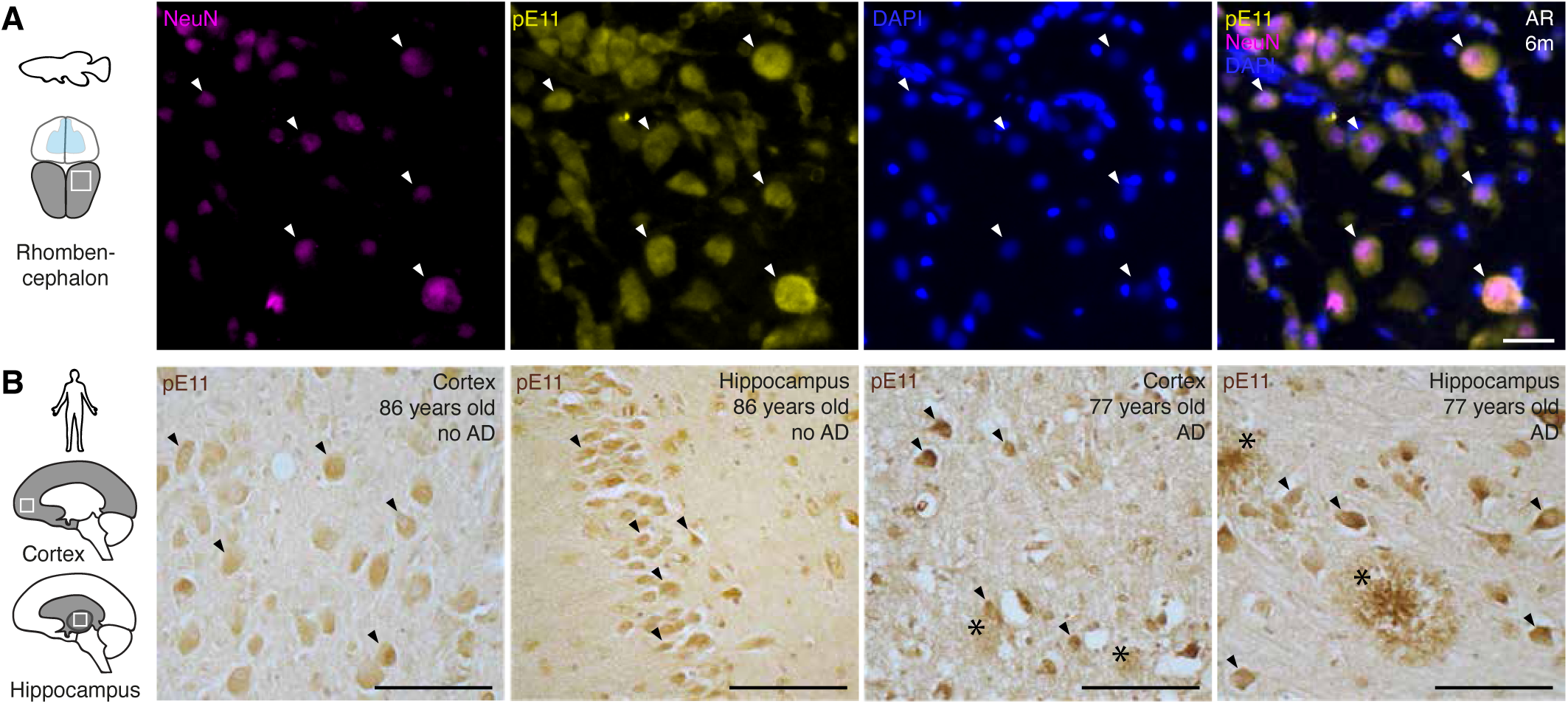
In aged killifish and human brains, pE11 signal predominantly localizes intraneuronally. (**A**) Immunohistochemistry staining in the 6-month-old anterior rhombencephalon (AR) using the pE11 antibody (yellow) and NeuN (Neuronal Nuclei) antibody. Arrowheads indicate colocalizing NeuN and pE11 signals. Scale bar represents 20 µm. (**B**) Immunohistochemistry using the pE11 antibody in two regions of the aged human brain of non-diseased (left) and Alzheimer’s diseased (right) brains. Arrowheads indicate intraneuronal signal, the asterisks indicate extracellular plaques. Scale bar represents 100 µm. Images are representative examples of three investigated individuals per condition.

**Figure S11.**
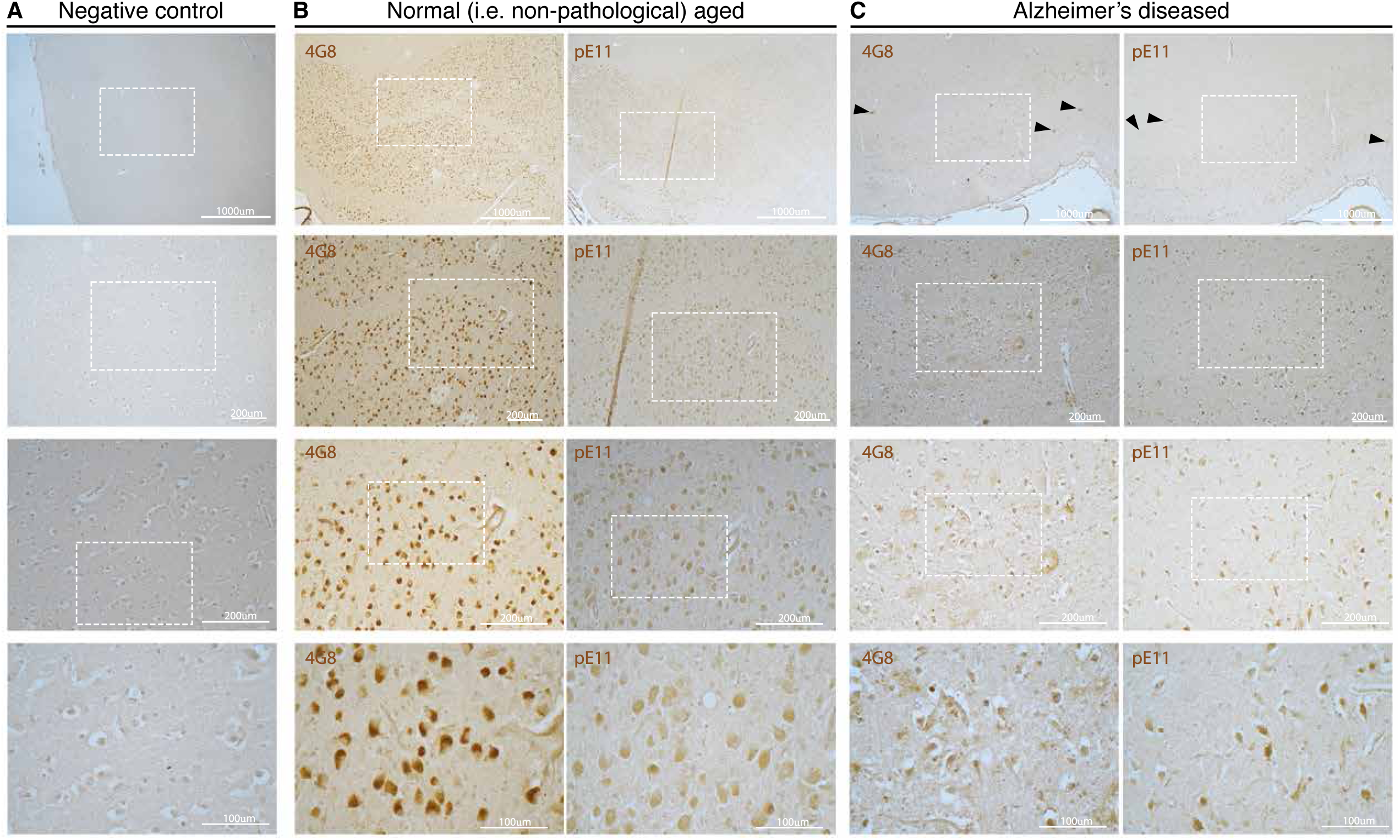
Staining of 4G8 and pE11 in human cortical areas. (**A**) Negative control staining for cortical human areas, realized without the addition of any primary antibody during the protocol. (**B**) Staining for 4G8 and pE11 in cortical areas of a normal (i.e. non-pathological) aged individual (>80 years old). (**C**) Staining for 4G8 and pE11 in cortical areas of an Alzheimer’s Disease diagnosed individual (>80 years old). (B, C) These images are representative examples of in total six investigated individuals. Arrowheads indicate extracellular amyloid plaques visualized by 4G8 (left) or pE11 (right).

**Figure S12.**
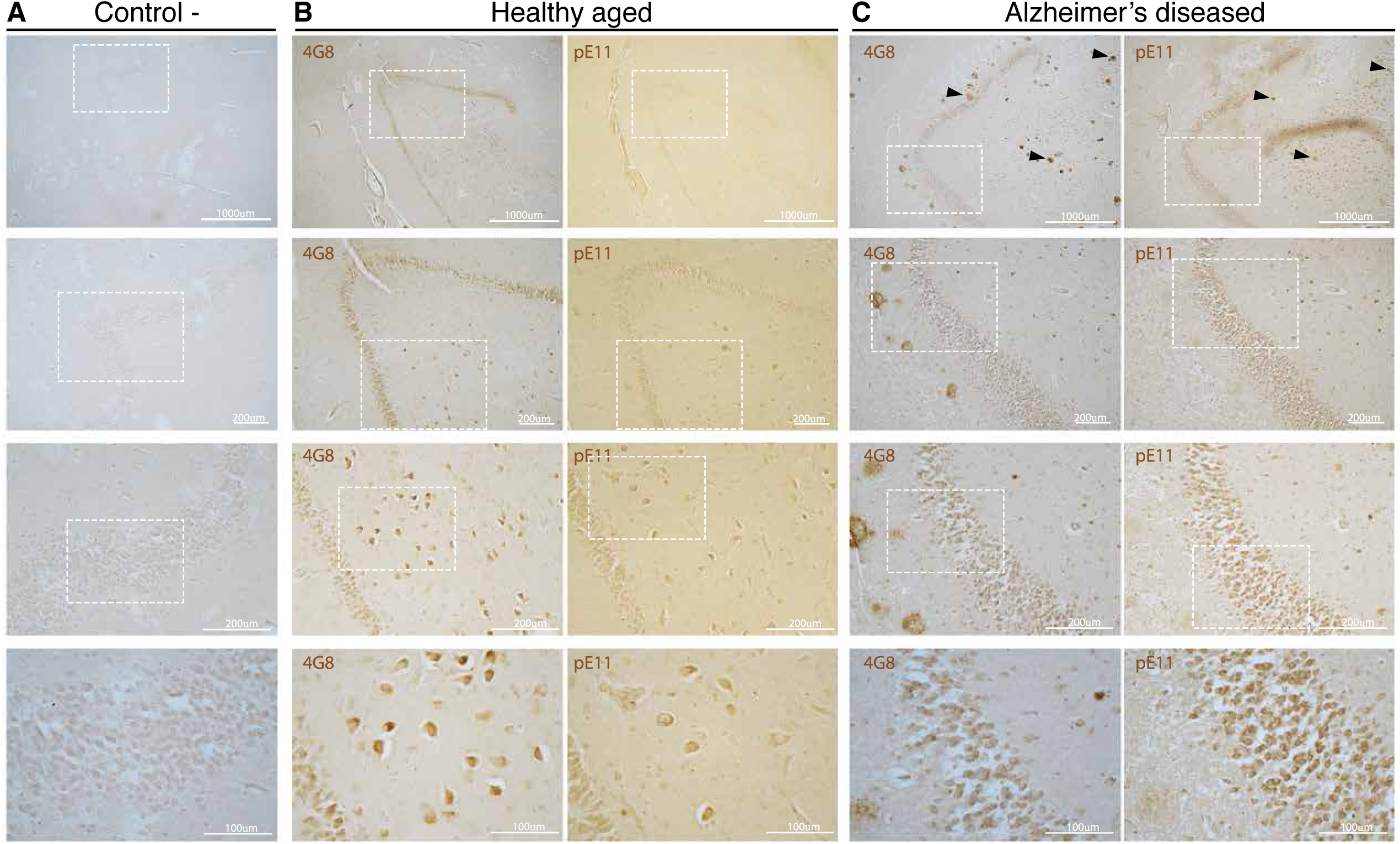
Staining of 4G8 and pE11 in human hippocampus. (**A**) Negative control staining for hippocampal human areas, realized without the addition of any primary antibody during the protocol. (**B**) Staining for 4G8 and pE11 in hippocampal areas of a control patient. (**C**) Staining for 4G8 and pE11 in hippocampal areas of an Alzheimer’s Disease affected patient. Arrowheads indicate extracellular amyloid plaques visualized by 4G8 (left) or pE11 (right).

**Figure S13.**
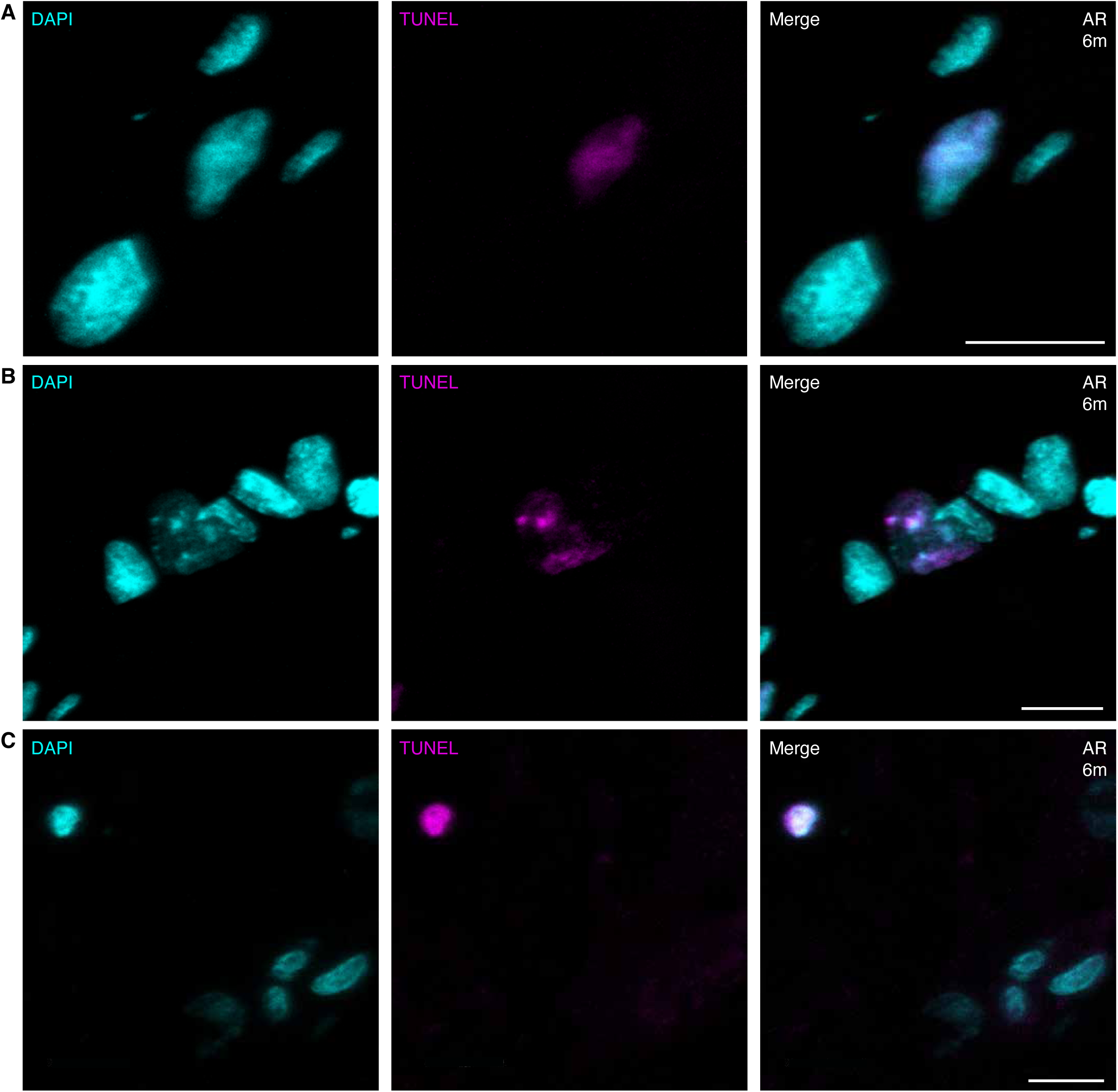
TUNEL staining in the anterior rhombencephalon. (**A-C**) Detection of apoptotic cells in the anterior rhombencephalon (AR) of a 6-month-old turquoise killifish through TUNEL staining. Various staining patterns were identified, including (**A**) visibly intact nuclei, (**B**) fragmented nuclei as well as (**C**) compacted nuclei. Scale bars represent 10 µm. (A-C) All three observed TUNEL staining patterns were observed in three biological replicates.

**Figure S14.**
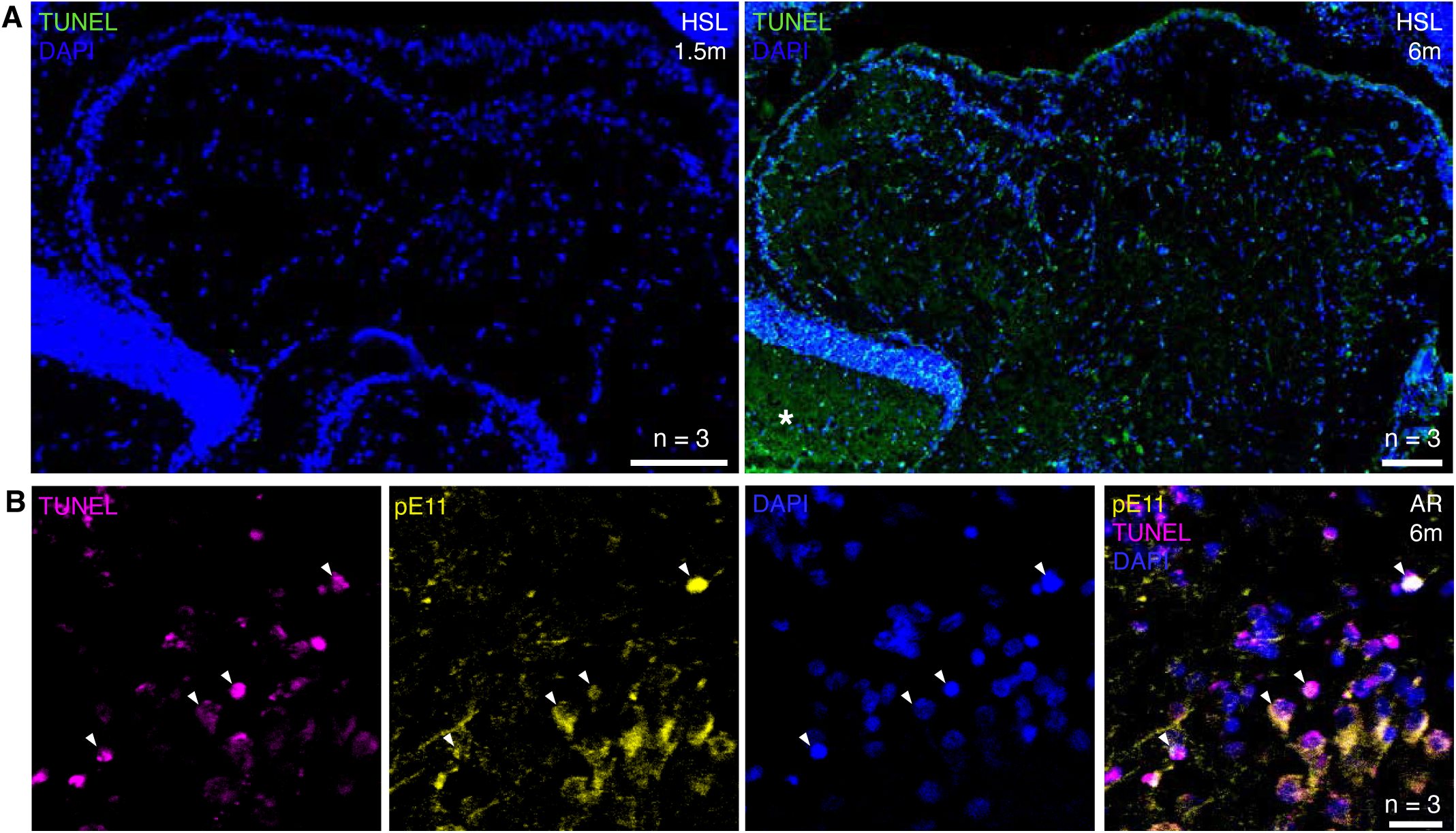
TUNEL staining shows a robust signal in aged turquoise killifish and overlaps with pE11 signal. (**A**) Detection of apoptotic cells in the hypothalamus superior lobe (HSL) of a young 1.5-month-old (**left**) or aged 6-month-old turquoise (**right**) turquoise killifish, visualized through TUNEL staining (green). The asterisk indicates age-related background signal, which was consistent throughout the apical tip of the optic tectum throughout the investigated brain sections of 6-month-old fish. Scale bars represent 100 µm. (**B**) Immunohistochemistry staining using the pE11 antibody (yellow) combined with TUNEL staining (magenta). Arrowheads indicate colocalizing signal. Scale bar represents 20 µm. AR = anterior rhomencephalon.

**Figure S15.**
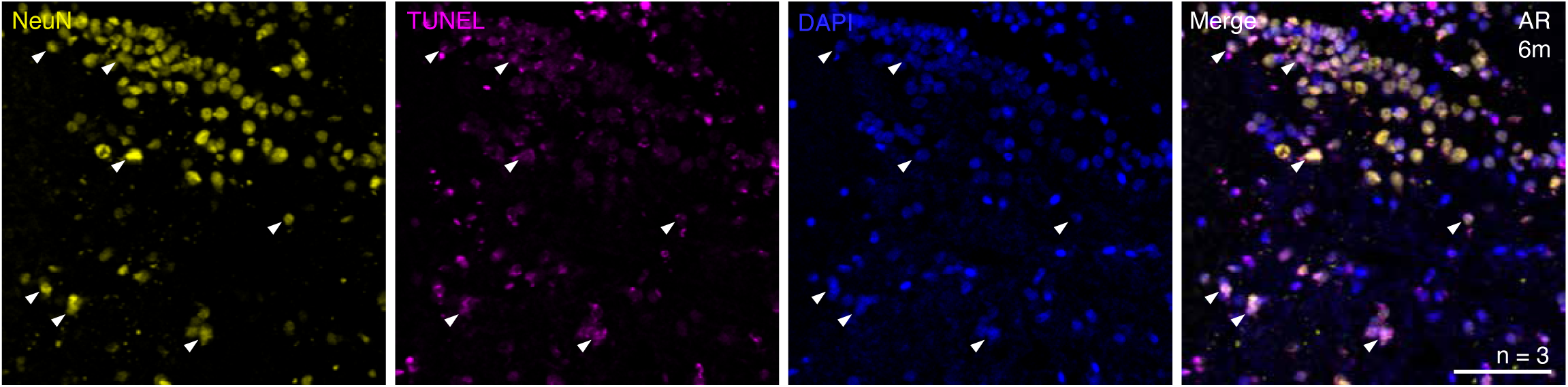
TUNEL staining colocalizes with NeuN in the aged anterior rhombencephalon of the turquoise killifish. (**A**) Immunohistochemistry staining using the pE11 antibody (yellow) combined with TUNEL staining (magenta). Arrowheads indicate colocalizing signal. Scale bar represents 20 µm. The staining was performed on the anterior rhombencephalon (AR) of a 6-month-old turquoise killifish and forms a representative example of three biological replicates.

**Figure S16.**
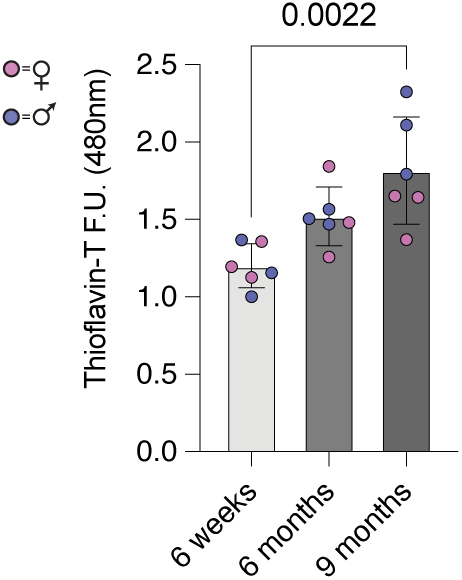
Beta-sheet aggregates accumulate in an age-dependent manner. Thioflavin-T assays were conducted on brain lysates of young, aged and old killifish. Shown values are 480nm emission levels 3 minutes after addition of Thioflavin-T to brain lysates. Values were normalized against the lowest measured value. Significance was calculated using a Mann-Whitney test.

**Figure S17.**
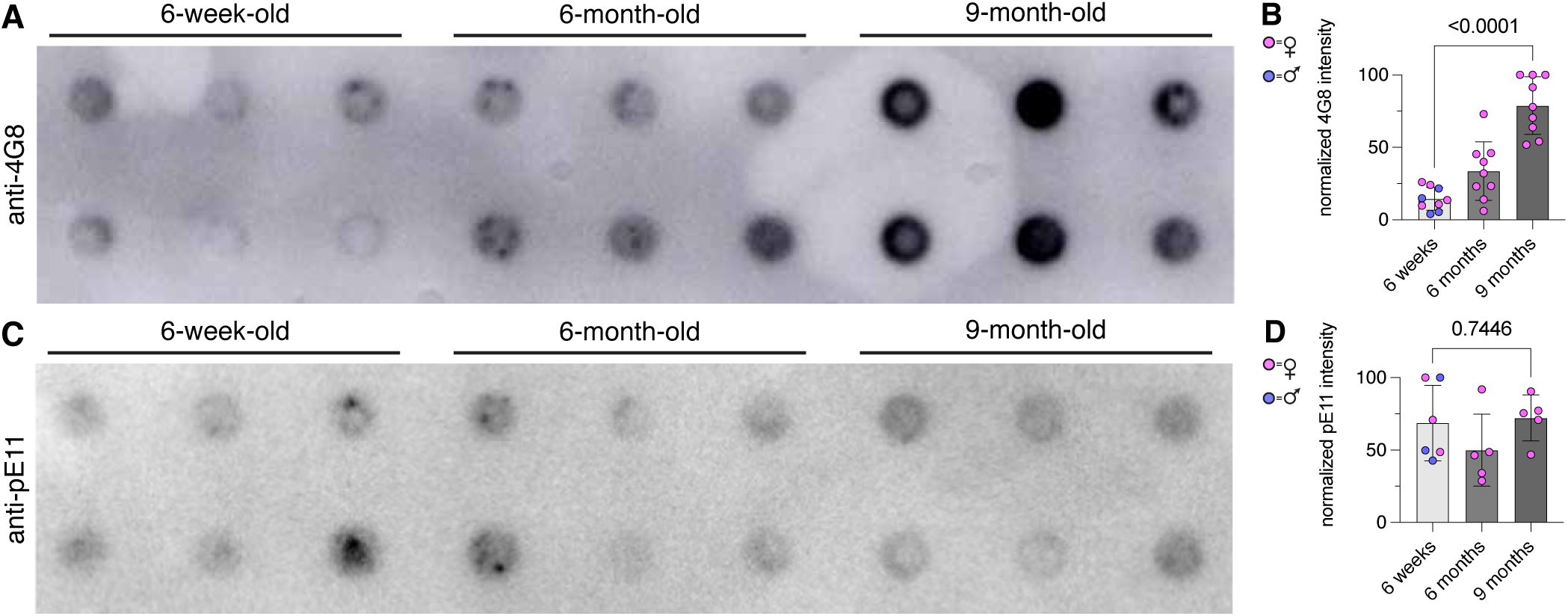
4G8+ peptides aggregate in the turquoise killifish brain in an age-dependent manner. (**A**) Filter retardation analysis of 4G8^+^ aggregates. 100 µg of protein from brain lysates from three individual killifish of each age group was spotted on the filter in duplicates (top/bottom) and probed with an anti-4G8 antibody. (**B**) Quantification of 4G8^+^ signal normalized against the maximum value measured per experiment. Each dot represents the averaged value of three technical replicates per brain. Error bars represent standard deviation. Significance was calculated using a Mann-Whitney test. (**C**) Filter retardation analysis of pE11^+^ aggregates. 100 µg of protein from brain lysates from three individual killifish of each age group was spotted on the filter in duplicates (top/bottom) and probed with an anti-pE11 antibody. (**D**) Quantification of pE11^+^ signal normalized against the maximum value measured per experiment. Each dot represents the averaged value of three technical replicates per brain. Error bars represent standard deviation. Significance was calculated using a Mann-Whitney test.

**Figure S18.**
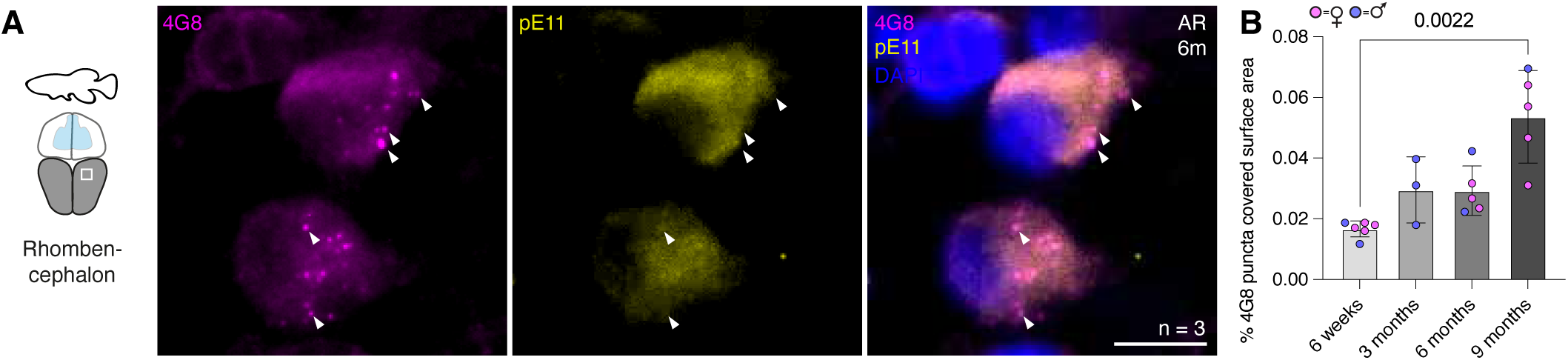
Intraneuronal 4G8^+^_/_pE11^-^ puncta increase with age in the turquoise killifish brain. (**A**) Immunohistochemistry staining in the 6-month-old anterior rhombencephalon (AR) using the pE11 antibody (yellow) and 4G8 antibody (magenta). Arrowheads indicate 4G8^+^/pE11^-^ puncta. Scale bar represents 10 µm. (**B**) Quantification of the percentage of 4G8^+^_-_puncta-covered surface area in AR regions at four different ages. Significance was calculated using a Mann-Whitney test. Error bars represent standard deviation.

**Figure S19.**
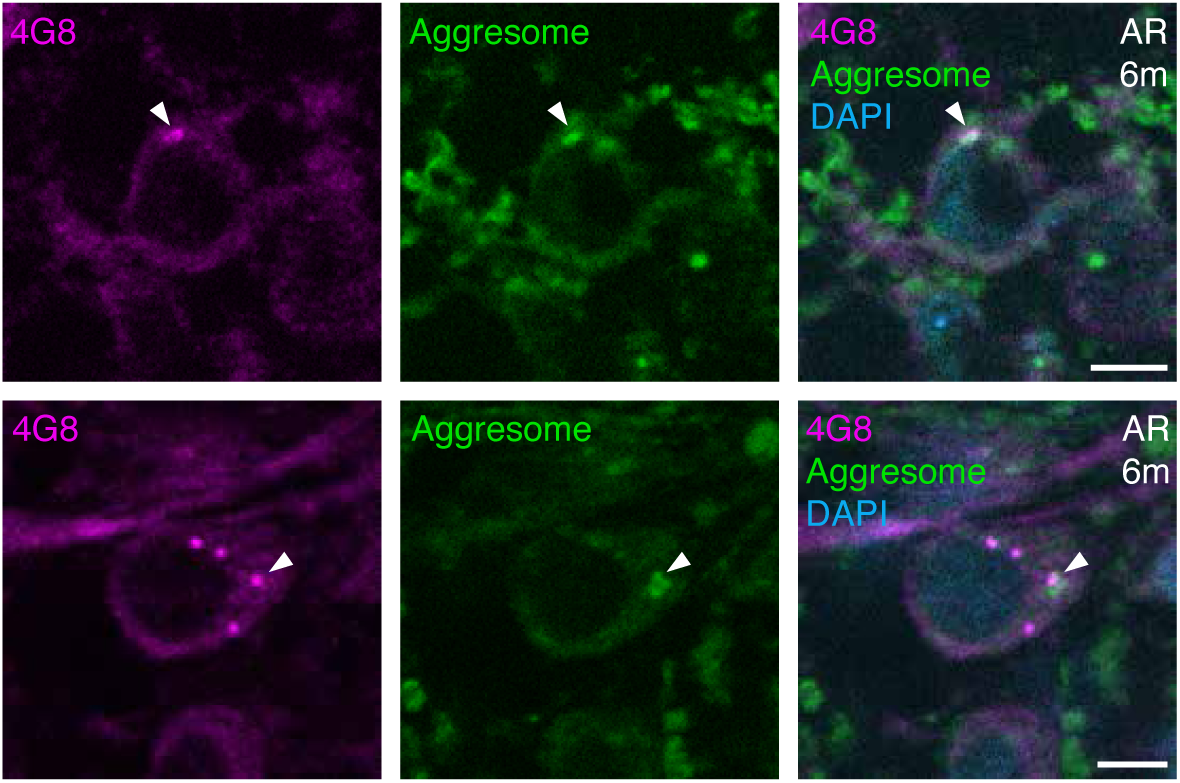
4G8 puncta partially overlap with Aggresome dye. Two examples of cells in the 6-month-old anterior rhombencephalon (AR) showing an overlap between the 4G8 antibody (magenta) and the PROTEOSTAT Aggresome dye indicating protein aggregates (green). Arrowheads indicate colocalization of Aggresome and 4G8 signal. Scale bars represents 5 µm.

**Figure S20.**
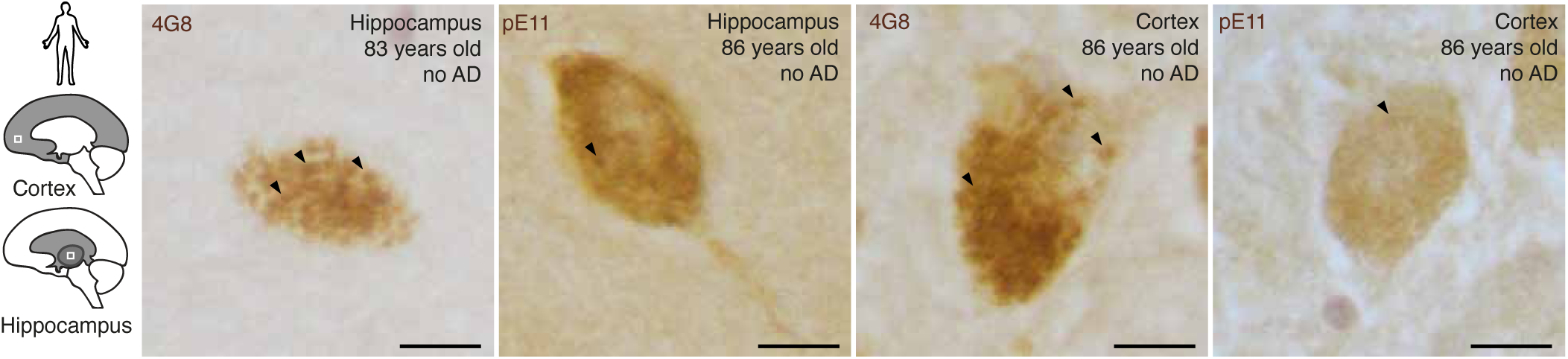
4G8 staining shows a granulated pattern, which is largely absent in the pE11 staining. Immunohistochemistry using the 4G8 and pE11 antibody in two regions of the non-diseased aged human brain. Arrowheads indicate intraneuronal puncta. Scale bar represents 10 µm.

**Figure S21.**
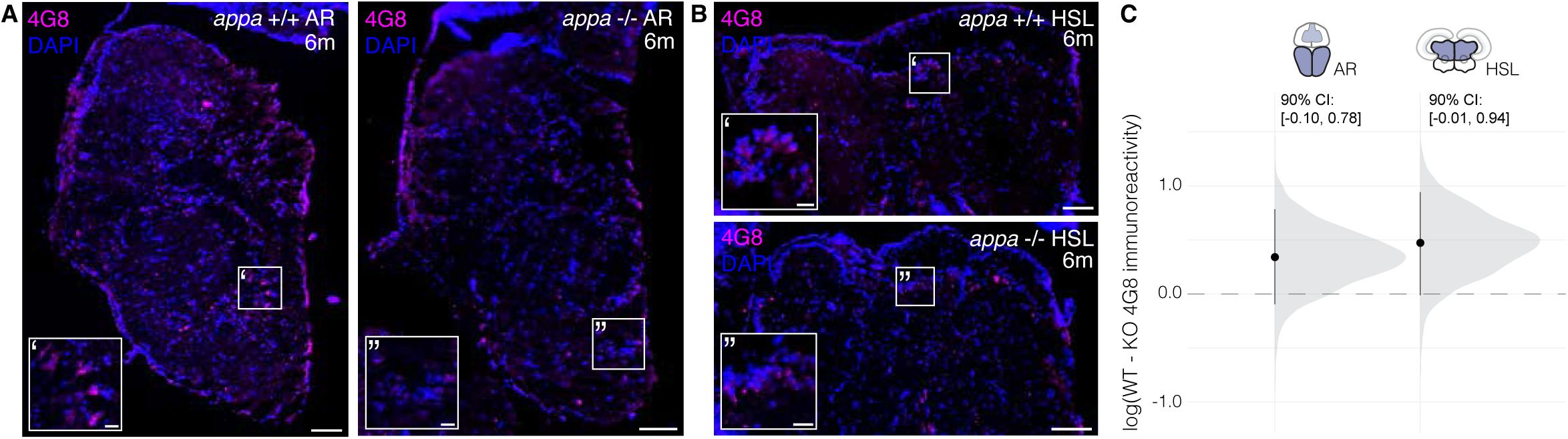
*appa* knock-out reduces 4G8 signal. (**A, B**) Representative immunohistochemistry staining results using the 4G8 antibody (magenta) in the anterior rhombencephalon (AR) of wild-type (A) and *appa* -/- mutant (B) brains. Scale bars represents 100 µm. (**C**) Posterior estimates of the genotype effect (WT − KO) on 4G8 immunoreactivity were obtained using a Bayesian model. Black vertical bars represent the 90% confidence intervals (CIs). The results indicate higher 4G8 immunoreactivity in *appa* wild-type (WT) brains compared to *appa* -/- (KO) brains in the anterior rhombencephalon (AR) and hypothalamus superior lobe (HSL) of 6-month-old killifish.

**Figure S22.**
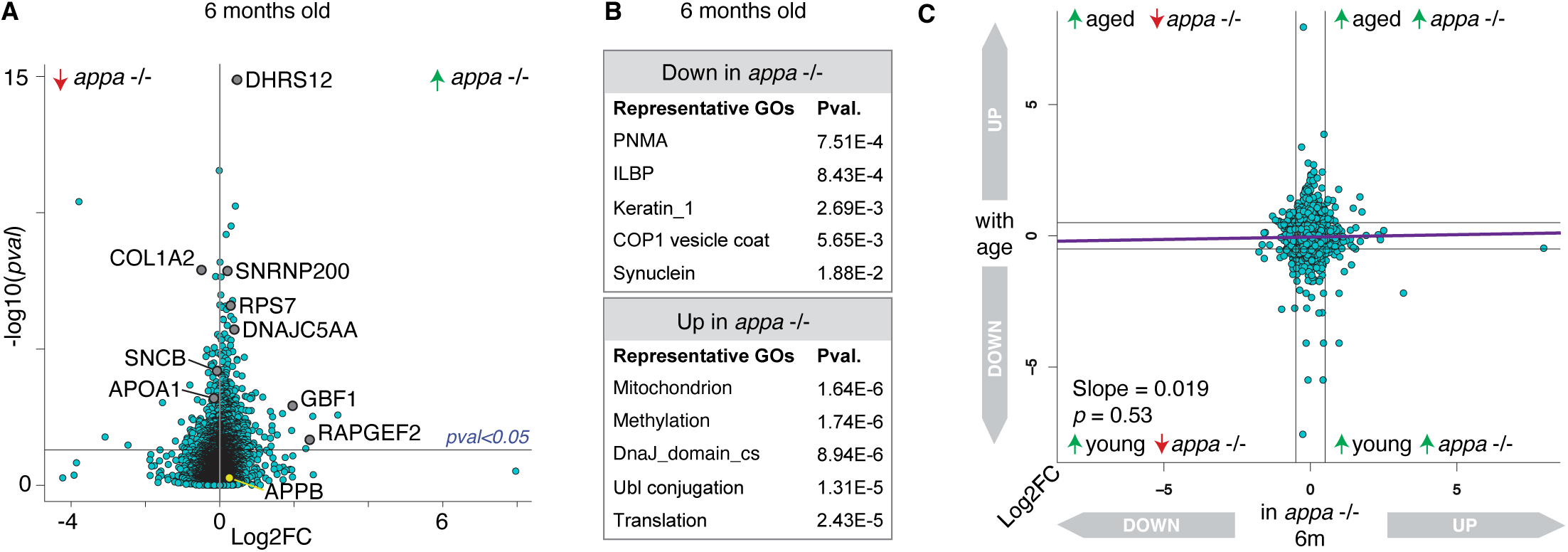
*appa* knock-out affects protein abundances at 6-months of age. (**A**) Volcano plot displaying differences in brain protein abundance between *appa* -/- mutants and their wild-type siblings (6-month-old, 3 males and 3 females per genotype). Differentially abundant proteins are highlighted, representing relevant biological processes. The vertical line indicates log2FC of 0. (**B**) Representative gene ontologies produced from the differentially abundant proteins between 6-month-old *appa* -/- mutants and their wild-type siblings (pval < 0.05). (**C**) Proteome-wide comparison between the log2FC of proteins up or downregulated in 6-months-old brains compared to 7-weeks-old brains (y-axis), plotted against the log2FC describing the up and downregulation of proteins in the *appa* mutant compared to the same aged wild-type siblings at 6 months.

**Figure S23.**
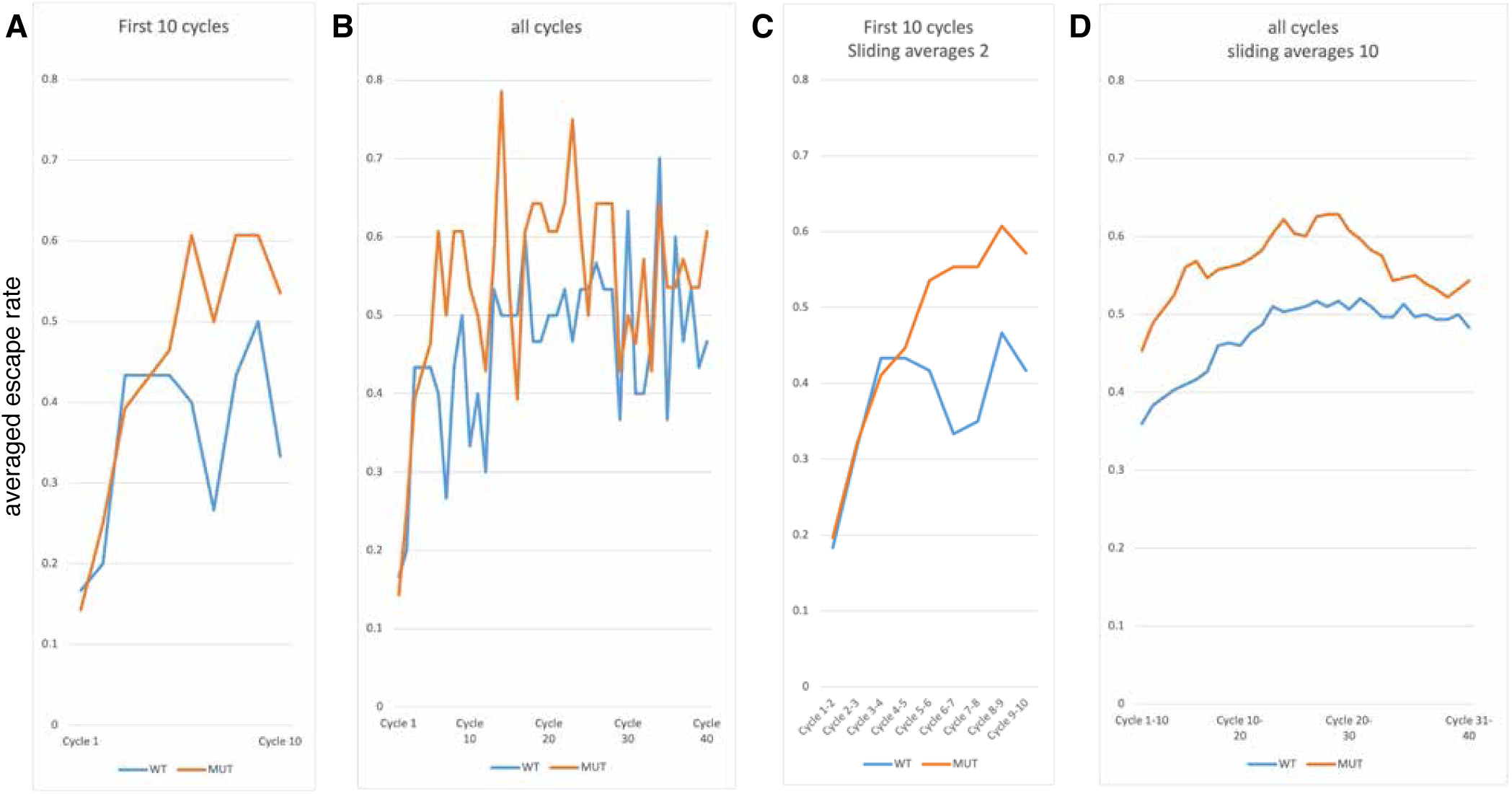
Averaged escape rates of *appa* -/- mutants and their wild-type siblings. (**A, B**) Plot describing the averaged escape rate of 7.5-month-old *appa* -/- mutants (orange, n = 28 fish) and their wild-type siblings (blue, n = 30 fish) across 10 (A) and the full 40 (B) cycles. (**C, D**) Sliding averaged values of the data presented in A and B, using either a sliding average of two (C) or ten (D) to increase the visibility of the general trends.

**Figure S24.**
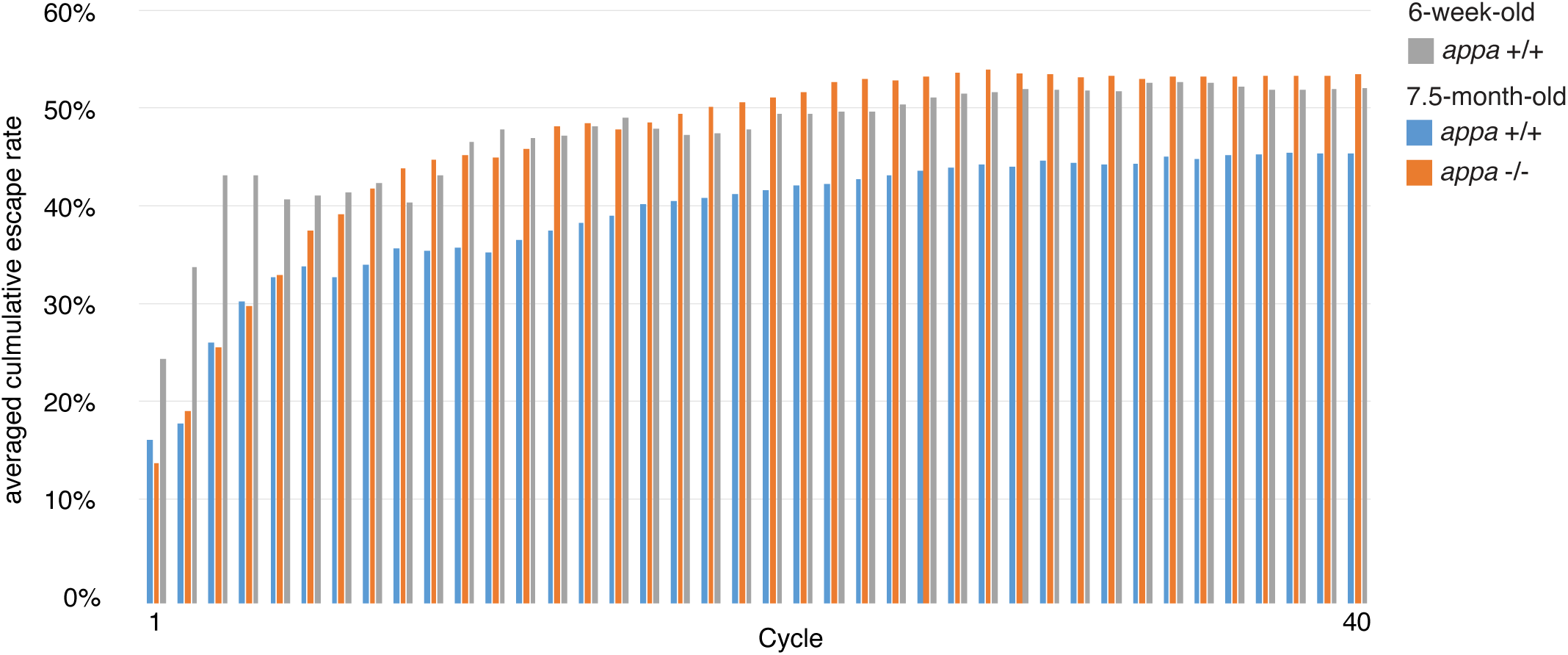
*appa* knock-out mitigates age-related learning deficiency in turquoise killifish. Plot describing the averaged cumulative escape rate of 7.5-month-old *appa* -/- mutants (orange, n = 28 fish) and their wild-type siblings (blue, n = 30 fish) across 40 cycles. In addition, a seperately conducted cohort of 6-week-old individuals has been tested and plotted (grey, n = 16 fish).

